# Temporal Metabolic Characteristics and Transcriptomic Landscape of Islets and Liver Reveal Dynamic Pathophysiology and Interorgan Crosstalk in High-fat Diet-induced Diabetes

**DOI:** 10.1101/2020.08.21.195453

**Authors:** Rui Gao, Qi Fu, He-Min Jiang, Min Shen, Rui-Ling Zhao, Yu Qian, Yun-Qiang He, Kuan-Feng Xu, Xin-Yu Xu, Heng Chen, Quan Zhang, Tao Yang

**Affiliations:** Department of Endocrinology and Metabolism, The First Affiliated Hospital of Nanjing Medical University, Nanjing, Jiangsu, 210029, China; Oxford Centre for Diabetes, Endocrinology and Metabolism, University of Oxford, Churchill Hospital, Oxford, OX37LE, UK

**Keywords:** High-fat diet-induced diabetes, Temporal transcriptomic analyses, Inter-tissue crosstalk, Islet dysfunction, Insulin resistance

## Abstract

**Objective:** Hyperinsulinemia and insulin resistance are co-existing characteristics of type 2 diabetes, whereas the molecular mechanism underlying this deleterious cycle remains elusive. The temporal transcriptomic landscape of core organs responsible for insulin secretion (islets) and insulin action (liver) could provide new insights.

**Methods:** The longitudinal profiling of glucose metabolism, insulin sensitivity, islet architecture and secretion were conducted in C57BL/6N mice fed a high-fat diet (HFD) or chow diet for 24 weeks. RNA-sequencing of islets and liver were performed once every 4 weeks. Weighted gene co-expression network analysis and Ingenuity Pathway Analysis were applied to construct networks and evaluate co-ordinated molecular interactions between islets and liver.

**Results:** Mice exhibited progressively deteriorated glucose homeostasis with hyperinsulinemia but impaired first-phase insulin secretion after 4 weeks on HFD. Insulin, glucagon and somatostatin secretion in response to glucose with or without palmitate gradually deteriorated from dysregulation to failure. Systemic insulin resistance developed over 24 weeks with variable time course in tissue-specific insulin action. Our transcriptomic datasets outlined the impact of HFD on dynamics of islet and liver molecular network at different stages. Correlation analyses revealed that both organs jointly programmed β-cell compensatory adaption via cell proliferation at early phase and irreversible islet dysfunction by inappropriate immune response at later stage. Alternations of T cell subpopulations validated the participation of adaptive immune response through priming and amplification phases in diabetic progression.

**Conclusion:** Our data provide a comprehensive landscape of crosstalk between islets and liver in diet-induced diabetes, elucidating the development of islet dysfunction and insulin resistance.

**Highlights:**

1. Diet-induced diabetes is featured by transition from islet dysfunction to failure
2. Insulin resistance develops with variable time course in different tissues
3. Dynamics of islet and liver molecular network interplay at different stages
4. Cell proliferation and improper immune reaction mediated interorgan crosstalk
5. Adaptive immune response participated via priming and amplification phases

## 1. INTRODUCTION

Over-nutrition is a major forerunner of type 2 diabetes mellitus (T2DM), which can both enhance secretion of insulin and attenuate its metabolic actions on peripheral tissues including liver, skeletal muscle and adipose tissue [1]. Hyperinsulinemia and insulin resistance are co-existing characteristics of T2DM, whereas which is the primary driver is still controversial. Mounting evidence over the decades proposed a paradigm that nutritional excess triggers peripheral tissue insensitivity to insulin, raising blood glucose level, which in turn promotes pancreatic β-cells to produce and secrete more insulin due to increase in demand [2,3]. However, recent observations proposed an opposite model that hyperinsulinemia is the causal factor for initiating insulin resistance in obesity-related diabetes [4,5]. These conflicting views indicated an incomplete understanding of the development of hyperinsulinemia and insulin resistance in T2DM, a key question regarding the aetiology of the disease.

Regardless of these contradictory perspectives, it is widely accepted that progressively deteriorated β-cell function and gradual loss of functional β-cell mass are the core events of diabetic development [6]. In order to identify molecules and mechanisms associated with β-cells’ transition from adaption to failure, “omics” technologies, particularly microarray-based transcriptomics and mass spectrometry-based proteomics, were broadly used on islets isolated from different T2DM animal models and human cadaver donors. However, despite over a decade of research, a comprehensive picture of diabetic progression is yet to be established. This is due to the fact that most studies: (1) focused on a single time point at relatively late stage of diabetes, which lacked temporal information of the disease progression [7,8]; (2) used rodent models that do not recapitulate the development of human diabetes, such as monogenic, polygenic or chemical-induced ones [8,9]; (3) lacked high-throughput sequencing data [10].

Individual β-cell can sense a multitude of signals and change its secretory responses according to metabolic demands. In recent years, endocrine and autocrine/paracrine factors have aroused great interest, however, a limited number of studies concentrated on the inter-organ crosstalk. Exogenous factors, such as humoral and neural signals originating from hepatocytes [11,12], adipocytes and various immune cells, not only constitute a significant link between obesity and insulin resistance, but also impact β-cell function and cell mass [13]. Liver, an important organ participating in fat metabolism, glycogen synthesis and decomposition, may play a role in regulating pancreatic β-cells. We previously identified that hepatocytes derived extracellular vesicles from high-fat diet (HFD) induced obese mice could modulate genes expression and promote proliferation of islet β-cells through miRNA [14]. During the last decade, other liver-derived circulating factors such as hepatic growth factor (HGF) [15,16], leukocyte-neutrophil elastase inhibitor (SerpinB1) [17], kisspeptin [18] and fibroblast growth factor 21 (FGF21) [19] have been reported to directly affect islet secretion, proliferation and regeneration.

In the present study, experiments were conducted to (1) define the timing of key physiological and molecular events in a rodent model of diet-induced diabetes characterized by hyperinsulinemia and insulin resistance; (2) determine the sequential repertoire of distinct mechanisms that underlie β-cells’ transition from dysfunction to failure; (3) assess the relationship of transcriptomic changes between islets (responsible for hyperinsulinemia) and liver (involved in insulin resistance), and determine the potential interconnected genes participating in this crosstalk. Thus, we established a dynamic profile of glucose metabolism, islet architecture and secretion, systemic and tissue-specific insulin resistance, and T cell subpopulations in diet-treated C57BL/6N mice over a time course of 24 weeks. Subsequent transcriptomic analyses of islets and liver unveiled the chronological order of T2DM-related molecular events during the deterioration of pancreatic islet function.

## 2. MATERIAL AND METHODS

### 2.1 Animals

Animal use and procedures were approved by the Medicine Animal Care Committee of Nanjing Medical University (IACUC1804001). All animal experiments were performed in accordance with the U.K. Animals (Scientific Procedures) Act, 1986 and associated guidelines. Four-week-old C57BL/6N male mice were purchased from GemPharmatech Co., Ltd. and were maintained at 22°C on a 12◻h light/dark cycle with *ad libitum* access to either chow diet (CD) (LabDiet 5001, 13.5% fat) or HFD (Research Diets D12492, 60% fat) for 4−24 weeks. Body weight, *ad libitum* blood glucose and food intake were monitored weekly. The average daily food intake was determined over 3 days by the net reduction in diet weight with the exclusion of spilled food. Energy intake is expressed as calories per mouse per day.

### 2.2 In vivo biochemical measurements

Serum insulin and glucagon levels were assessed using the Mouse Insulin and Glucagon ELISA kits (Mercodia, Uppsala, Sweden) according to the manufacturer’s instructions, respectively. For the measurement of glucagon, ethylenediaminetetraacetic acid and aprotinin were placed in sample collection tubes beforehand. For other biochemical tests, blood was collected from the retro-orbital sinus after overnight fast. Serum alanine aminotransferase (ALT), aspartate aminotransferase (AST), alkaline phosphatase (ALP), albumin (ALB), total cholesterol (T-CHO), triglyceride (TG), high-density lipoprotein cholesterol (HDL-C), and low-density lipoprotein cholesterol (LDL-C) levels after 16 h fasting were determined using an automatic biochemical analyzer (HITACHI 7100, Hitachi Koki Co. Ltd., Hitachinaka City, Japan).

### 2.3 Glucose and insulin tolerance tests

Intraperitoneal glucose tolerance tests (IPGTTs) and insulin tolerance tests (ITTs) were respectively performed after food withdrawal for 16 h or 5 h at intervals of 4 weeks. Tail blood glucose was measured (Medisafe Mini, Terumo, Japan) at baseline and periodically following intraperitoneal administration of 2 g/kg D-glucose for IPGTT or 0.75 U/kg insulin (Humulin R, Eli Lilly, Indianapolis, IN, USA) for ITT. At 5, 15, 30, 60 and 120 min after glucose injection during IPGTT, additional blood (25 μL) was collected via the tail vein for subsequent insulin determination.

### 2.4 Hyperinsulinemic-euglycemic clamps

Mice were subjected to carotid artery and jugular vein catheterization for sampling and infusion, respectively. The catheters were kept patent with heparin sodium (1 IU/mL) and body weights were recorded daily during recovery. After mice were fasted for 5 h, the clamp started at t=0 min (end of the 5 h fast) with a continuous insulin infusion (Humulin R, Eli Lilly, Indianapolis, IN) at a rate of 4 mU/kg/min. Glucose infusion rate (GIR) was adjusted based on the measurement of blood glucose at 5 min intervals to maintain euglycemia at 6.1−7.2 mmol/L (110−130 mg/dL) during the steady-state period from t=80−120 min. Plasma insulin levels were determined from blood samples obtained from the carotid artery at t=-5 min prior to the insulin infusion and t=120 min. Insulin sensitivity index (ISI) is defined as ISI_Clamp_=GIR/(GxΔI) where GIR is normalized for G (steady-state blood glucose concentration) and ΔI (difference between fasting and steady-state plasma insulin concentrations) [20].

### 2.5 Purification of primary mouse islets

After clamped with a hemostat at the duodenal opening, the bile duct was cannulated and perfused with 2◻mL collagenase P (Roche Diagnostics, Germany, 1◻mg/mL in Hanks’ balanced salt solution (HBSS)). Pancreatic tissue was digested at 37◻°C for 10◻min. The reaction was stopped by adding cold HBSS with 10% fetal bovine serum (FBS), followed by 10Lseconds of vigorous shaking. Islets were separated onto a gradient composed of HBSS and Histopaque (11191 and 10771, Sigma Aldrich, USA) layers and were hand-picked for 2 times under a dissection microscope to minimize acinar cells. The purity of islets for RNA-sequencing was evaluated by the expression of *Amy1a* (a marker of acinar cells) and *Krt19* (a marker of ductal cells). In our transcriptomic dataset, *Amy1a* and *Krt19* showed no statistical difference at mRNA level from any time points, which indicates the contaminants were fairly minimized.

### 2.6 In vitro hormone secretion measurements

Islets were isolated and pooled from 4−5 mice per group, then incubated overnight in RPMI-1640 supplemented with 10% FBS and 1% Pen/Strep. The next day, groups of 20 size-matched islets were pre-incubated for 1 h in Krebs-Ringers-Bicarbonate-HEPES (KRBH) buffer, containing (in mmol/L): 140 NaCl, 3.6 KCl, 2 NaHCO_3_, 0.5 NaH_2_PO_4_, 0.5 MgSO_4_, 5 HEPES, 2.6 CaCl_2_, 1 glucose and 0.2% (wt/vol.) BSA (pH 7.4 with NaOH). Following this, islets were sequentially subjected to KRBH solutions containing 2 or 20 mmol/L glucose with or without 0.5 mM palmitate (PA) for 1 h at 37°C in 5% CO_2_. PA was non-covalently conjugated to fatty acid-free BSA at a 6.6:1 molar ratio. Supernatants were collected and islets were lysed overnight in acid ethanol buffer (75% EtOH, 0.55% HCl). Insulin (Mercodia, Uppsala, Sweden), glucagon (Mercodia, Uppsala, Sweden) and somatostatin (Phoenix Pharmaceuticals, CA, USA) in supernatants and contents were determined using the ELISA kits following the manufacturer’s instructions. Secreted hormones were calculated as percentage of total contents per hour.

### 2.7 Electron microscopy

For transmission electron microscopy (TEM), isolated islets from 3 mice per group were pooled and cultured overnight. After pre-incubation and treatment with 2 or 20 mM glucose in KRBH buffer for 1 h at 37°C, islets were fixed in 2.5% glutaraldehyde and 2.5% paraformaldehyde in cacodylate buffer (0.1 M, pH 7.4). Then, samples were post-fixed in 1.0% osmium tetroxide for 2 h and stained with 2% (wt/vol.) uranyl acetate in double distilled water to increase the contrast. After washing and dehydration through graded alcohol (50, 70, 90 and 100%), samples were embedded in EPON 812, and ultrathin sections (60−80 nm thick) were cut by a diamond knife using a Leica ultra-microtome EM UC6 (Leica, Wetzlar, Germany). Islets were examined by FEI Tecnai G2 Spirit Bio TWIN (FEI, Hillsboro, USA) with an accelerating voltage of 80 kV. Mature insulin granules were determined and quantified by grayscale threshold analyses. Immature granules, as well as docked granules, were identified and counted manually. Granules located within 100 nm from the plasma membrane and without signs of fusion were considered docked [21].

### 2.8 Histopathological and immunohistochemical staining

After the mice were transcardically perfused with 4% (wt/vol.) paraformaldehyde, pancreas and livers were dissected from each group, fixed in 4% paraformaldehyde and dehydrated in a graded sucrose series. The paraffin-embedded pancreatic samples were then continuously sectioned at 5 μm, rehydrated with a sequential wash in xylene, 100%, 95%, 75% ethanol and water, and then boiled in 1 mM ethylenediaminetetraacetic acid for antigen retrieval. After blocking with 5% goat serum, slides were incubated overnight with primary antibodies: anti-insulin (ab7842, 1:400), anti-glucagon (ab10988, 1:400), anti-somatostatin (ab30788, 1:100), and anti-Ki67 (ab15580, 1:1000) (Abcam, Cambridge, UK). The next day, sections were washed and incubated with conjugated or biotinylated secondary antibodies for 1 h at room temperature. The paraffin-embedded liver tissues were also sliced at 5 μm, and stained with haematoxylin and eosin (H&E) for liver structure, Masson’s Trichrome and Sirius Rose for collagen fiber deposition according to the standard protocols (Solarbio, Beijing, China). Primary antibody anti-CD31 (ab28364, 1:50) (Abcam, Cambridge, UK) was used for immunostaining to assess hepatic microvessel density. Images were captured using Axiovert A1 Inverted Microscope (Carl Zeiss, Jena, Germany).

### 2.9 Western blot

Liver, quadriceps femoris, gastrocnemius and white adipose tissue were harvested 10 min after an intraperitoneal bolus of insulin (5 U/kg) or PBS from overnight fasted mice. After homogenized in RIPA buffer (R0278, Sigma Aldrich, USA) with 1:100 dilution of protease inhibitor cocktail (P8340, Sigma Aldrich, USA), tissue extracts were centrifuged at high speed (15,000 g) to eliminate insoluble material. Protein concentrations in the supernatants were measured using the bicinchoninic acid assay (Pierce BCA Protein Assay Reagent, Thermo Scientific, Waltham, MA, USA). Equal amount of protein lysates (40 mg) was separated by SDS-polyacrylamide gel electrophoresis (SDS-PAGE). Proteins were then transferred to PVDF membranes (Millipore, Burlington, MA, USA) and blotted with anti-phospho-Akt (Ser473) (4060, 1:1000), anti-Akt (4691, 1:1000), anti-phospho-GSK-3α/β (Ser21/9) (8566, 1:1000), anti-GSK-3α/β (5676, 1:1000) and anti-β-Actin (13E5, 1:1000) (Cell Signaling, Danvers, MA, USA). The western blot procedure was performed as previously described. Values were expressed as phospho-protein over total protein.

### 2.10 Flow cytometry

Anti-mouse CD3 fluorescein isothiocyanate (FITC) (100306), anti-mouse CD4 FITC (100510), anti-mouse CD8 peridinin-chlorophyll protein (PerCP)-cyanine 5.5 (Cy5.5) (100734), anti-mouse CD19 PerCP-Cy5.5 (115534), anti-mouse CD25 allophycocyanin (APC) (101910), anti-mouse inducible costimulator (ICOS) phycoerythrin (PE)-cyanine 7 (Cy7) (313520), anti-mouse C-X-C motif chemokine receptor 5 (CXCR5) Brilliant Violet 421^TM^ (145511), anti-mouse interferon (IFN)-γ PE (505808), anti-mouse interleukin (IL)-4 PE (504104) and anti-mouse IL-17 PE (517008) (BioLegend, San Diego, CA, USA) were used. For forkhead box P3 (Foxp3) intracellular staining, anti-mouse Foxp3 PE and the Foxp3 Fix/Perm Kit (eBioscience, San Diego, CA, USA) were used according to the manufacturer’s instructions.

Mouse single cell suspensions of spleens, draining lymph nodes (including peri-pancreas lymph nodes) were prepared. Cells were first stained with surface antibodies for 30 min at 4°C in the dark. After surface staining, cells were fixed, permeabilized for 40 min in 1×Foxp3 Fix/Perm Buffer and washed twice with 1× Perm Buffer. Cells were then incubated with Foxp3 antibody for 30 min. For intracellular cytokines detection, cells were stimulated with phorbol myristate acetate (PMA) and ionomycin in the presence of brefeldin A (BioLegend, San Diego, CA, USA) for 5 h before surface staining. Cells were then fixed with Intracellular Fixation Buffer (BD Biosciences, San Diego, CA, USA), incubated with Permeabilization Buffer (BD Biosciences, San Diego, CA, USA), and then stained with anti-mouse IFN-γ PE, anti-mouse IL-4 PE or anti-mouse IL-17 PE. The cytometric data were collected using a FACS Aria II Sorp flow cytometer (BD Biosciences, San Diego, CA, USA).

### 2.11 RNA extraction, library construction and sequencing

Total RNA was isolated from handpicked islets and homogenized liver tissue using TRIzol reagent (Invitrogen, CA, USA) in accordance with the manufacturer’s instructions. The quality and quantity of extracted RNA were measured by on-chip electrophoresis utilizing the Agilent RNA 6000 Nano Kit and Agilent 2100 Bioanalyzer (Agilent Technologies, CA, USA). Samples exhibited 1.9≤A260/A280◻≤◻2.2, RNA integrity number>8.0 and 28S/18S>1.0 were selected for library preparation. The total RNA was purified by enriching poly (A) mRNA with magnetic oligo (dT) beads. After reverse transcription by using the random hexamer (N6) primers, double-stranded cDNA fragments were synthesized and subjected to end repair and adapter ligation. The cDNA libraries were then constructed by PCR amplification, and finally sequenced on BGISEQ-500 sequencing platform by the Beijing Genomics Institute (Shenzhen, China).

### 2.12 Reads filtering and de novo assembly

The sequencing raw reads were filtered for low-quality, adaptor-polluted, high content of unknown base reads by SOAPnuke (v1.5.2) [22]. We used Trinity (v2.0.6) [23] to perform de novo assembly, and Tgicl (v2.0.6) [24] on cluster transcripts to remove redundancy and get unigenes. The high-quality clean reads were then mapped to the mouse reference genome (GRCm38) via HISAT2 (v2.0.4) [25] and full-length transcriptomic database via Bowtie2 (v2.2.5) [26] (**Supplemental Table 1**). The gene expression levels were then quantified by RSEM (v1.1.12) [27] and were normalized by the method of fragments per kilobase of exon model per million reads mapped (FPKM). The DEseq2 (fold change≥2 and adjusted *P*-value≤0.05) package was applied to identify differentially expressed genes (DEGs) between HFD and CD groups at each time point.

### 2.13 Pathway analyses

To interpret the functional significance of DEGs, Ingenuity Pathway Analysis (IPA) (Qiagen, Redwood City, CA, USA, www.qiagen.com/ingenuity) was conducted to determine enriched canonical pathways, upstream regulators, mechanistic networks, and diseases and functions. DEGs with a log_2_fold change (log_2_FC) either ≤−1 for down-regulated or ≥1 for up-regulated, adjusted *P*-value ≤0.05 and average FPKM >1 were selected. Pathway significance is expressed as *P*-value calculated by right-tailed Fisher’s exact test, which indicates the possibility that the correlation between DEGs from our dataset and a given process/function is due to random chance. Kyoto Encyclopedia Genes and Genomes (KEGG) analyses of DEGs were also performed according to the gene annotations provided by DAVID online tool (v6.8).

Secretome data were obtained from Vertebrate Secretome Database (VerSeDa) [28], which stores information about proteins that are predicted to be secreted through the classical and non-classical mechanisms, for a wide range of vertebrate species deposited at NCBI, UCSC and ENSEMBL sites.

### 2.14 Weighted gene co-expression networks between islets, liver and metabolic traits

We separately conducted weighted gene co-expression network analyses (WGCNA) for 48 islet and 48 liver samples. Gene modules (clusters of genes displaying similar correlated patterns of transcription) were built by employing the WGCNA package in R [29], following the general guidelines [30]. Two inclusion criteria for filtering genes in WGCNA construction were applied: (1) genes with FPKM>0 in any of the 48 samples were selected; (2) median absolute deviation (MAD) was calculated for each gene as a robust measure of variability, and the 75% most varying genes were chosen. Briefly, pairwise Pearson correlation coefficients between all included genes were calculated to generate a signed similarity. After a sensitivity analysis of scale-free topology (R^2^>0.9), soft threshold power was respectively set to 6 for islets and 18 for liver to obtain a weighted adjacency matrix. Then it converted into a topological overlapping matrix network which was used as input for a hierarchical clustering analysis. The modules were finally identified by implementing the dynamic hierarchical tree-cut algorithm, using the parameters deepSplit=2 and minClusterSize=20. Module eigengenes which were defined as the principal components of each module were calculated using the moduleEigengenes function.

We associated the islet and liver modules by measuring the Pearson correlation using the WGCNA “relating modules to external information” analyses. The significance (Pearson correlation) was calculated for each gene in islet modules to the corresponding correlated liver modules, and for each gene in liver modules to the corresponding correlated islet modules. We also obtained the connectivity (membership) of each gene to its own module. Pairwise comparisons of all modules could identify a set of ‘key genes’ which were most correlated to the modules of other tissue (>0.5) and also displayed a high membership in its own module (>0.9).

To identify gene modules associated with traits such as fasting glucose level, we also performed the same analyses (“relating modules to external information”) to correlate islet and liver modules to metabolic variables. The cut-off of *P*-value for selecting modules of interest was set to 0.05. Modules of interest were further characterized by KEGG pathway enrichment analyses.

### 2.15 Quantitative real-time PCR

To validate the reliability of data obtained by RNA-sequencing, quantitative real-time PCR (qRT-PCR) was performed. Total RNA was extracted and its quality and quantity were examined as indicated above. 1 μg of total RNA was reverse transcribed using Takara PrimeScript™ RT Master Mix (Clontech Laboratories, USA). qRT-PCR was performed on Step One Plus Real-Time PCR System (Applied Biosystems, USA) with SYBR Premix Ex Taq II Kit (Clontech Laboratories, USA) and primers presented in **Supplemental Table 2**. β-Actin was served as an internal control and relative changes in mRNA expression were calculated by the comparative ΔCt method.

### 2.16 Statistical analysis

The images presented in the manuscript are representative of the data and the image/staining quality. The area occupied by glucagon or somatostatin staining (area A) was measured using Image J and expressed relative to that of insulin staining (area B) by the formula A / (A+B). For quantification of β-cell proliferation, nuclear Ki67^+^ Ins^+^ cells were counted as proliferating β-cells. The percentage was calculated by dividing the number of Ki67^+^ Ins^+^ cells by the total number of Ins^+^ cells in each islet. All values are expressed as mean±standard error of mean (SEM) and compared by Student’s unpaired *t*-test or analysis of variance (ANOVA) with Tukey’s test which corrects for multiple hypotheses. Correlation was quantified by the Pearson’s r for parametric data analysis. Area under the curve (AUC) was calculated by trapezoid analysis and was compared by *t*-test. A two-sided *P*-value<0.05 was considered statistically significant. Graphs and statistical analyses were completed using GraphPad Prism 6.0 (GraphPad Software, San Diego, CA, USA).

### 2.17 Data availability

The transcriptomic datasets generated and analyzed during the current study have been deposited in Gene Expression Omnibus (GEO) repository with the accession code GSE153222. All other data that support the findings of this study are available from the corresponding author upon reasonable request.

## 3. RESULTS

### 3.1 Experimental design and general metabolic characteristics of HFD in C57BL/6N mice

To illustrate the global characteristics and molecular dynamics in HFD model, we monitored the temporal profile of glucose metabolism, islet architecture and secretion, and tissue-specific insulin sensitivity in C57BL/6N mice fed on a 60% HFD or a CD for 24 weeks. Transcriptomes of islets and liver at six consecutive time points with the interval of 4 weeks (week 4, 8, 12, 16, 20 and 24) were analyzed using WGCNA and IPA (**Figure 1A**). As to non-fasting morning blood glucose level, we identified mild hyperglycemia after 1 week of HFD. From week 8 to week 14, largest difference in non-fasting plasma glucose was observed. Notably after week 20, although insignificant, plasma glucose in HFD remained ~1 mmol/L higher than that of CD (**Figure 1C**). There was also a modest increase in energy consumption in HFD mice compared with CD ones during the study period (**Figure 1D**). Interestingly, caloric excess was disproportionately stored in adipose tissue throughout 24 weeks in HFD group (**Supplemental Figure 1A**). The biochemical measurements in **Supplemental Figure 1B** presented HFD-induced hepatoxicity as revealed by increase in serum liver enzymes and aminotransferases, and HFD-induced dyslipidemia as shown by elevation in cholesterol levels. In concordance with impaired liver function, we also identified gradually deteriorated hepatic steatosis, cytoplasmatic ballooning, fibrosis and angiogenesis in HFD mice (**Supplemental Figure 2A**).

**Figure 1:**
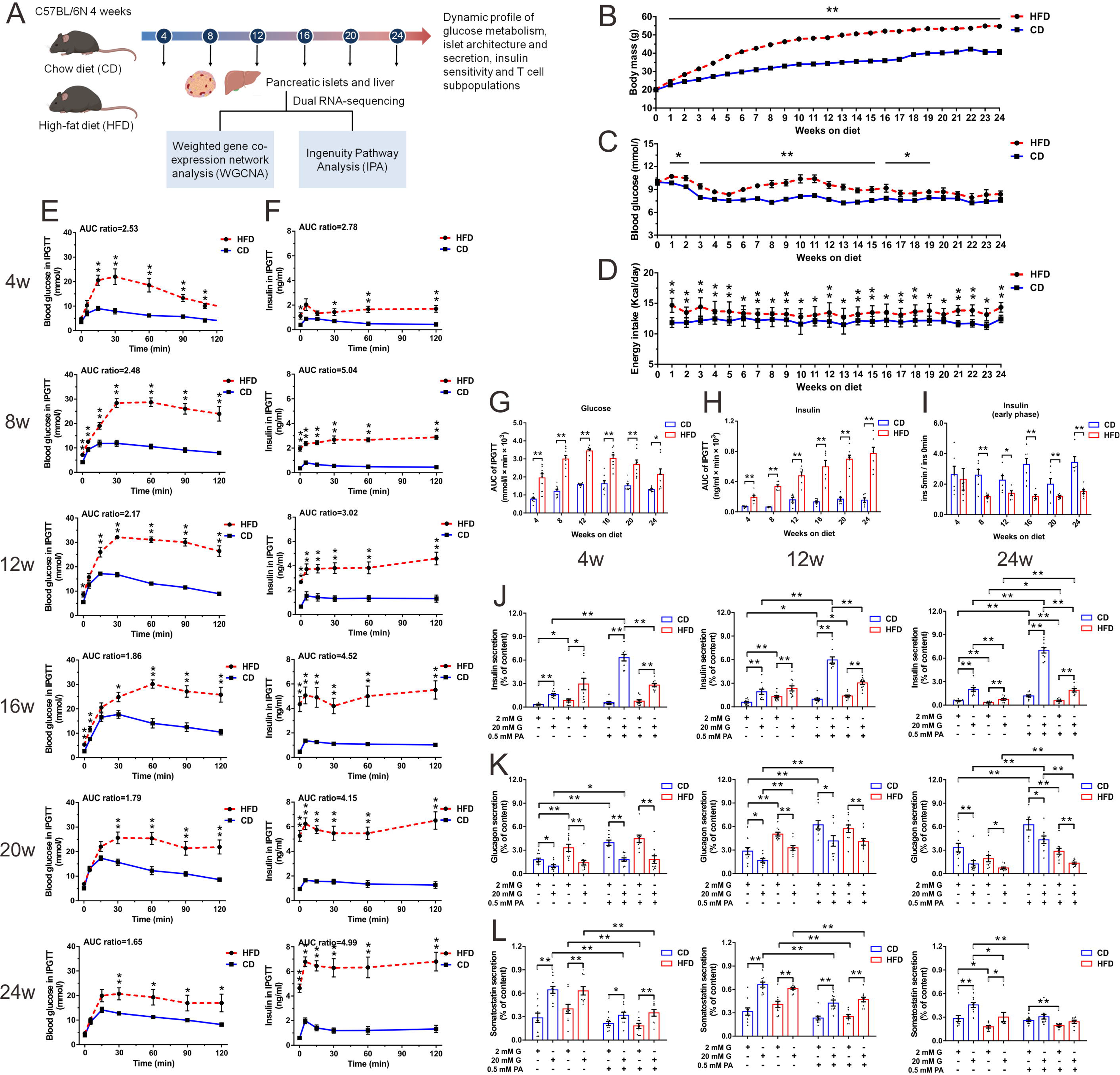
Progressively impaired glucose homeostasis and islet dysfunction of HFD mice both in vivo and in vitro. **(A)** Schematic of the experimental design. **(B, C, D)** Weekly monitored body weight (N≥16 mice/group) (B), morning ad libitum blood glucose (N≥14 mice/group) (C) and caloric intake (N≥4 cages/group) (D). **(E, F)** Blood glucose (E) and the corresponding plasma insulin concentration (F) during IPGTT (2 g/kg glucose) (N≥6 mice/group). The ratios of AUC for HFD vs. CD are presented at the top of each graph. **(G, H)** AUC calculation for glucose (G) and insulin (H) during IPGTT. **(I)** First-phase insulin secretion defect in HFD mice indicated by fold change of insulin level at 5 min with respect to 0 min. **(J, K, L)** Insulin (J), glucagon (K) and somatostatin (L) secretion in islets isolated from HFD and CD mice after 4, 12 and 24 weeks of diet (N=4−5 mice/group). All data are expressed as mean±SEM and analyzed using unpaired two-tailed *t*-test. \**P*<0.05, \*\**P*<0.01.

### 3.2 Progressively impaired glucose homeostasis and islet function of HFD mice both in vivo and in vitro

IPGTTs were conducted every 4 weeks to monitor islet function during 24 weeks of diet treatments. Compared with CD mice, HFD mice developed glucose intolerance as early as 4 weeks, presenting with a significantly higher peak and slower decay of plasma glucose following glucose challenge (**Figure 1E**). As a result, the integrated glucose level (AUC) in HFD was around 2-fold of that seen in CD for the entire duration of 24 weeks (**Figure 1G**). However, this glucose intolerance slightly ameliorated after 16 weeks of HFD and fasting hyperglycameia attenuated at week 20, which may be attributable to the compensatory increase in β-cell function and/or mass in response to the diet. Indeed, the corresponding insulin level revealed a progressively exaggerated and significantly delayed glucose-induced secretory response in HFD mice (**Figure 1F**). The integrated insulin secretion (AUC) during glucose challenge increased from 2.78-fold (week 4) to 4.99-fold (week 24) of that seen in CD group (**Figure 1H**). Notably, we observed an evidently elevated fasting insulin level in HFD mice at week 4, prior to the development of fasting hyperglycaemia at week 8. Despite the overall enhanced insulin secretion, the first phase glucose-induced insulin secretion (insulin level at 5 min relative to basal release) was impaired from week 8 on HFD (**Figure 1I**). Fasted and 2 h refed plasma glucagon levels suggested α-cell dysfunction and subsequently inappropriate glucagon release were not responsible for hyperglycaemia in HFD mice (**Supplemental Figure 1C**).

We also measured insulin, glucagon and somatostatin secretion in intact islets from mice on HFD or CD at week 4, 12 (with the highest blood glucose level during IPGTT) and 24 (with the highest serum insulin level during IPGTT) in response to low and high glucose in the presence or absence of PA. The fold increase in insulin secretion stimulated by 20 mM glucose alone (3.53-fold in HFD vs. 4.77-fold in CD at week 4; 1.81-fold in HFD vs. 3.25-fold in CD at week 12; and 2.29-fold in HFD vs. 3.62-fold in CD at week 24) or 0.5 mM PA alone (0.90-fold in HFD vs. 1.72-fold in CD at week 4; 1.07-fold in HFD vs. 1.65-fold in CD at week 12; and 1.79-fold in HFD vs. 2.12-fold in CD at week 24) was both modestly reduced. PA potentiated glucose stimulated insulin secretion (GSIS) was also impaired at these three time points (3.72-fold in HFD vs. 11.03-fold in CD at week 4, 2.20-fold in HFD vs. 6.12-fold in CD at week 12, and 3.34-fold in HFD vs. 5.95-fold in CD at week 24). It is noticeable that islets from 4 weeks and 12 weeks of HFD exhibited a higher basal insulin release (2 mM glucose in the absence of PA) in comparison with CD. However, after 24 weeks of treatment, insulin secretion of HFD mouse islets was significantly lower than CD with or without the presence of stimulatory glucose or PA concentration, indicating β-cell function failure (**Figure 1J**). Glucagon secretion of both HFD and CD mouse islets showed intact secretory response to glucose and PA at all three time points. Similar to that of insulin, a more robust basal release of glucagon was observed in HFD at week 4 and 12, albeit with an evident decrease at week 24 (**Figure 1K**). Somatostatin secretion in HFD mice displayed marginally decreased sensitivity in response to the stimulatory action of 20 mM glucose and the inhibitory action of PA at 20 mM glucose compared with CD. Similar to insulin and glucagon secretion, an elevated basal somatostatin release was only observed in HFD islets at week 4 and 12, followed by a significantly reduced secretion at week 24 (**Figure 1L**). Altogether, these results demonstrated specific hyposensitivity of HFD islets to glucose and PA despite enhanced basal insulin, glucagon and somatostatin secretion at early stage.

### 3.3 Dynamic changes in islet morphology, cell composition and ultrastructure

Immunofluorescent staining was performed to examine the changes in islet size distribution and cell composition at different time points of diet treatments. In comparison with CD mice, the islet size did not change significantly after HFD feeding for 4 weeks, whereas at week 12, the prevalence of large islet population (6000−7999 μm^2^) doubled and that of small islets (<2000 μm^2^) relatively decreased in HFD group. The difference was even more profound after 24 weeks of HFD (**Figure 2B**). The glucagon positive area ratio of HFD islets began to decline from week 16 despite of concomitant augmentation in islet size (**Figure 2C**). The somatostatin positive area ratio of HFD islets also decreased by nearly 2-fold at week 8 and continued to reduce over 24 weeks of feeding period (**Figure 2D**). Pancreatic immunohistochemical examination corroborated the findings of progressively enlarged islet mass and increased abundance of Ins^+^ β-cells in HFD mice (**Supplemental Figure 2B**). To test whether the hyperplasia of β-cells was due to increased proliferation, we next quantitated proliferation markers such as Ki67 in β-cells at different time points of diet treatments. In CD mice, Ki67^+^ Ins^+^ β-cells were barely detectable. The percentage of Ki67^+^ Ins^+^ β-cells was only transiently increased in HFD mice at week 4 and 8, and showed no significant difference to CD during the subsequent weeks (**Figure 2F**).

**Figure 2:**
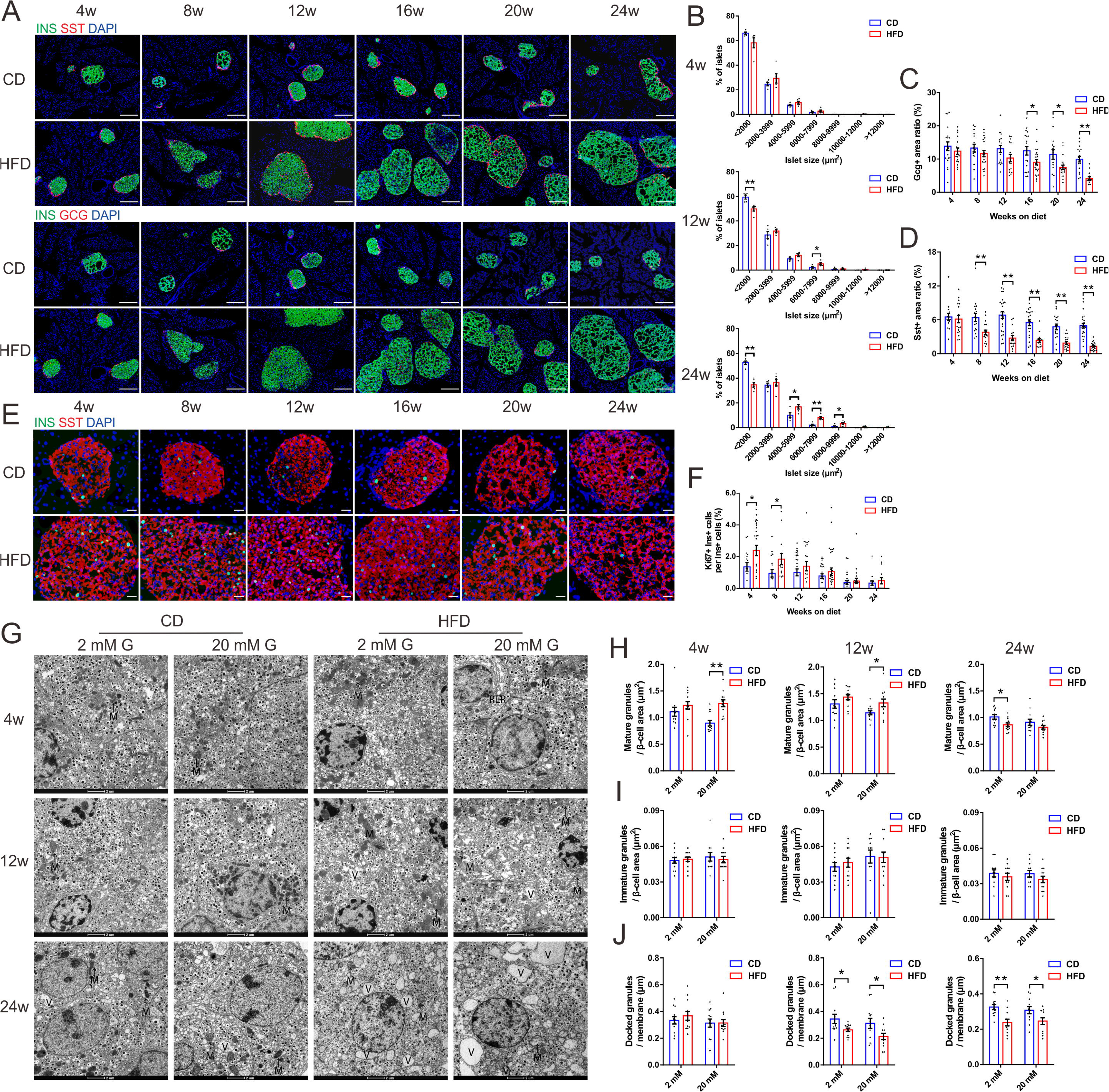
Dynamic changes in islet morphology, cell composition and ultrastructure. **(A)** Representitive immunofluorescent images showing consecutive pancreatic sections double-labeled either for insulin and glucagon or insulin and somatostatin (scale bar: 50 μm). **(B)** Islet size distributions analyzed by morphometry (N=5 mice/group). **(C, D)** Quantification of glucagon (C) and somatostatin (D) stained area (N=15−25 islets/group, N=4−5 mice/group). **(E)** Representitive immunofluorescent images showing Ki67^+^ Ins^+^ cells in pancreatic sections (scale bar: 10 μm). **(F)** Quantification of the percentage of Ki67^+^ Ins^+^ cells in total Ins^+^ cells (N≥14 islets/group, N=3−5 mice/group). **(G)** Representitive electron micrographs showing β-cells with insulin-containing granules after 2 mM or 20 mM glucose stimulation. Mitochondria (M), endoplasmic reticulum (RER) and vacuoles (V) are marked (scale bar: 2 μm). **(H, I, J)** Quantification of mature (H), immature (I) and docked (J) insulin granules in β-cells of HFD and CD islets (N=12 β-cells/group, N=6−8 islets/group, N=3 mice/group). All data are expressed as mean±SEM and analyzed using unpaired two-tailed *t*-test. \**P*<0.05, \*\**P*<0.01.

We next analyzed the ultrastructure of β-cells from HFD and CD mice by TEM. At week 4 and 12 of the diet, there was no significant difference in density of mature granules after 2 mM glucose treatment between two groups. However, 20 mM glucose stimulation led to an evident increase in mature granule density in HFD. Interestingly, in β-cells of mice fed on HFD for 24 weeks, the density of mature granules in 2 mM glucose was significantly reduced compared with CD. This supported the view from in vitro GSIS results that there was a transition from enhanced to impaired insulin secretion in HFD mice (**Figure 2H**). With regard to immature granules, no significant difference was observed (**Figure 2I**). We also calculated the density of docked granules and identified that HFD resulted in a different distribution of granules with fewer granules docked at the cell membrane compared to CD (**Figure 2J**). This may explain the reduced first phase of glucose-induced insulin secretion in HFD mice.

β-cells from HFD mice also showed several ultrastructural alterations (**Figure 2G**). HFD β-cell mitochondria were round-shaped rather than elongated, with fragmented cristae, reduced electron density and augmented volume. There was also massive accumulation of vacuoles characterized by the presence of closed membranes surrounding organelles and cytoplasmic portions in β-cells, possibly suggesting dysregulated autophagy. Interestingly, α-cells appeared ultrastructurally normal and well-granulated, whereas δ-cells were characterized by degranulation in HFD-treated group (**Supplemental Figure 2C**), which was quite similar with the TEM features observed in T2DM patients [31].

### 3.4 Longitudinal assessment of systemic and tissue-specific insulin sensitivity in HFD mice

To assess whether the impaired glucose tolerance in HFD was due to defects in insulin sensitivity, longitudinal ITTs were performed with an interval of 4 weeks. Compared with CD mice, HFD mice presented a relatively intact insulin-induced hypoglycemic response at week 4, while overt insulin resistance was only detectable after 8 weeks of diet treatment (**Figure 3A**). Hyperinsulinemic-euglycemic clamps (HI/EG) were also conducted to evaluate systemic insulin sensitivity. At week 4, HFD mice had a non-evident decrease in GIR compared with CD (ISI=0.90±0.08 in HFD vs. ISI=1.17±0.10 in CD at week 4, *P*=0.060). However, after 12 and 24 weeks of treatment, a significant lower GIR was required for HFD mice to maintain euglycemia at steady state (ISI=0.42±0.14 in HFD vs. ISI=1.79±0.43 in CD at week 12, *P*=0.013; ISI=0.15±0.04 in HFD vs. ISI=0.98±0.16 in CD at week 24, *P*<0.001) (**Figure 3B**).

**Figure 3:**
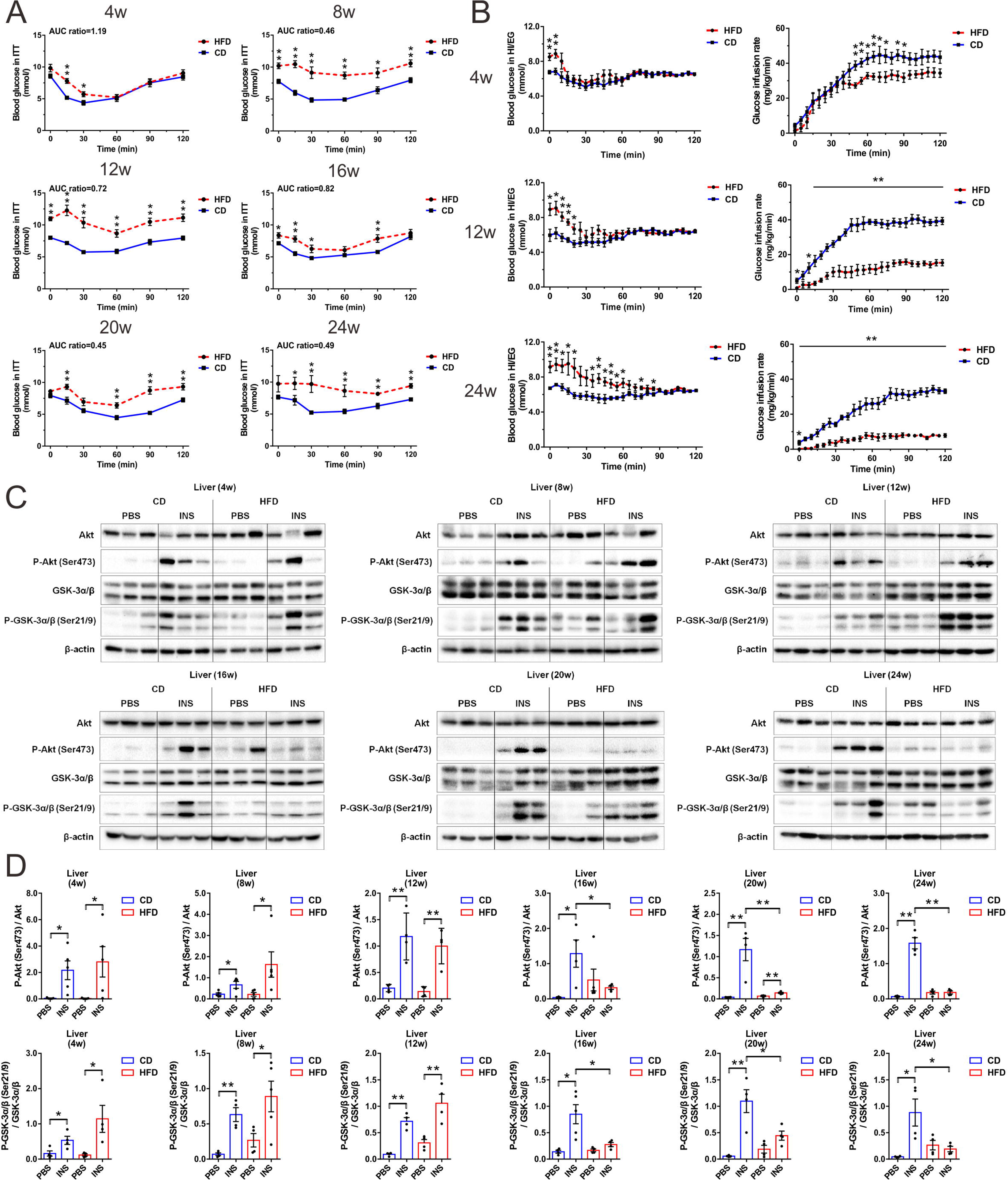
Longitudinal assessment of systemic and tissue-specific insulin sensitivity in CD and HFD mice. **(A)** Blood glucose during ITT (0.75 U/kg insulin) (N≥7 mice/group). The ratios of AUC for HFD vs. CD are presented at the top of each graph. **(B)** Hyperinsulinemic-euglycemic clamp tests (N≥6 mice/group). **(C, D)** Immunoblot (C) and quantification (D) of Akt (Ser473) phosphorylation status relative to total Akt and GSK-3α/β (Ser21/9) phosphorylation status relative to total GSK-3α/β in liver (N=4−6 mice/group). All data are expressed as mean±SEM and analyzed using unpaired two-tailed *t*-test. \**P*<0.05, \*\**P*<0.01.

For tissue-specific insulin action as displayed in **Figure 3C and D**, the hepatic acute response to insulin was not different between HFD and CD before week 16. However, phosphorylation of Akt after insulin injection was significantly lowered by 75.46% (week 16), 87.85% (week 20), and 88.34% (week 24) afterwards. The phosphorylation of GSK-3α/β in liver also demonstrated similar patterns. As regard to quadriceps femoris (**Supplemental Figure 3A and B**) and gastrocnemius tissue (**Supplemental Figure 3C and D**), the insulin-stimulated increases in p-Akt and p-GSK-3α/β were also blunted in HFD starting from week 16. Of note, in adipose tissue (**Supplemental Figure 3E and F**), reduced insulin response was evident since week 8 on HFD, prior to that in liver and skeletal muscle. This phenomenon might be attributed to adipose tissue being the primary site of nutrient storage. These data revealed that the development of insulin resistance at different organs had variable time course, with liver and skeletal muscle initiating from week 16 and adipose tissue starting from week 8.

### 3.5 Transcriptomic profiles and pathway dynamics of islets and liver during diabetes progression

To unveil the molecular mechanisms underlying these metabolic changes during diabetes progression, RNA-sequencing and transcriptomic analyses of islets and liver were respectively performed in quadruplicates at six consecutive time points of diet treatments (week 4, 8, 12, 16, 20 and 24). In total, 3844 DEGs were found in islets, of which 33 were shared among all six time points (**Figure 4C**). With regard to liver, 4101 DEGs were discovered throughout 24 weeks of feeding, of which 39 were overlapped (**Figure 4D**). To validate the transcriptomic results, 10 DEGs were randomly selected, and their expression patterns measured by qRT-PCR in independent HFD/CD islet and liver samples were very similar to those in RNA-sequencing data (**Supplemental Figure 4A** and **Supplemental Figure 4B**).

**Figure 4:**
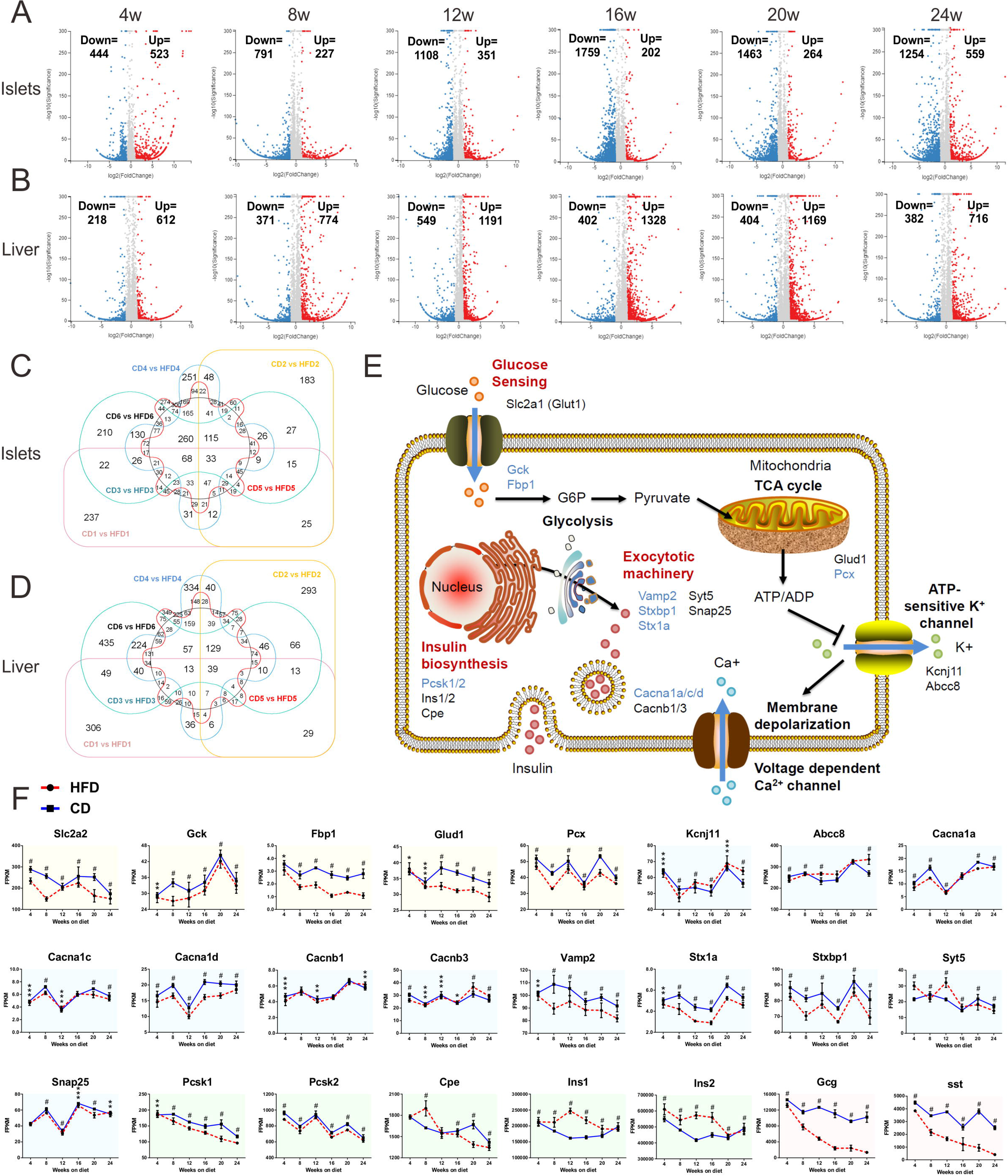
Transcriptomic profiles and pathway dynamics of islets and liver during diabetes progression. **(A, B)** Islet (A) and liver (B) volcano plots. The numbers of up-regulated (red dots) and down-regulated (blue dots) genes are marked in each graph. **(C, D)** Islet (C) and liver (D) venn diagrams of identifiable and quantifiable DEGs. **(E)** Schematic representation illustrating the physiology and key genes of GSIS in pancreatic β-cells. Genes that were significantly down-regulated are colored in blue. **(F)** The time course expression data for genes involved in GSIS. Yellow, blue, green and red shaded areas respectively refer to genes participating in glucose sensing, exocytotic machinery, insulin biosynthesis and the secretion of other islet cells. All data are expressed as mean±SEM. *adjusted *P*<0.05, **adjusted *P*<0.01, ***adjusted *P*<0.001, ^#^adjusted *P*<0.0001.

For pathway dynamics analyses, we investigated the temporal patterns with representative genes involved in islet hormone secretion, pancreatic endocrine cell development and hepatic insulin resistance. **Figure 4E and F** depicted the primary signal flow of GSIS in pancreatic β-cells, showing some down-regulated genes in glucose sensing and metabolism (*Gck and Fbp1*), Ca^2+^ flux (*Cacna1d*), granule docking and release (*Vamp2, Stxbp1 and Stx1a*). The pathway of insulin biosynthesis (*Cpe* and *Ins1*) was up-regulated before week 16 while gradually down-regulated afterwards, which might indicate a transition from compensatory oversecretion to functional impairment. We also concluded the gene expression of a synopsis of factors regulating endocrine pancreas development over time (**Supplemental Figure 5**). Certain genes participating in the differentiation of pancreatic progenitors into endocrine progenitors (*Gata6*, *Sox9* and *Onecut1*) were down-regulated throughout 24 weeks. Furthermore, **Supplemental Figure 6** provided a schematic representation of mechanisms involved in hepatic insulin resistance. Pathways of lipid-induced insulin resistance (PKC∊ (*Prkce*) and PP2A (*Sptlc1/2* and *Ppp2ca*)), intracellular inflammatory signaling (TLR4 (*Tlr4*) and IKK (*Ikbkb*)), and unfolded protein response (IRE1α (*Ern1*), BiP (*Hspa5*) and PERK (*Eif2ak3*)) were significantly up-regulated in HFD mice.

KEGG enrichment (**Figure 5A and B**) and IPA (**Figure 5C and D**) of DEGs were carried out to demonstrate top over-represented canonical pathways in each tissue. In islets, the early perturbations were characterized by up-regulation of cell replication pathways (cell cycle and oocyte meiosis in KEGG; cell cycle: G2/M DNA damage checkpoint regulation in IPA), which was coincident with the assessment of proliferative antigens in histological staining. Another notable feature is the gradual up-regulation in glycolysis/gluconeogenesis since week 4 on HFD, probably suggesting that defect in oxidative phosphorylation would lead to the anaerobic metabolism being dominant in producing energy. During week 16−24, signaling pathways associated with adaptive immune responses were significantly enriched in islets (T cell receptor signaling pathway, Th1 and Th2 cell differentiation, and Th17 cell differentiation in KEGG; B cell development, and altered T cell and B cell signaling in rheumatoid arthritis in IPA). It is also noticeable that some enriched pathways in liver were correlated with islet function and diabetic progression, such as insulin secretion, type I and type II diabetes, maturity onset diabetes of the young, and AGE-RAGE signaling pathway in diabetic complications.

**Figure 5:**
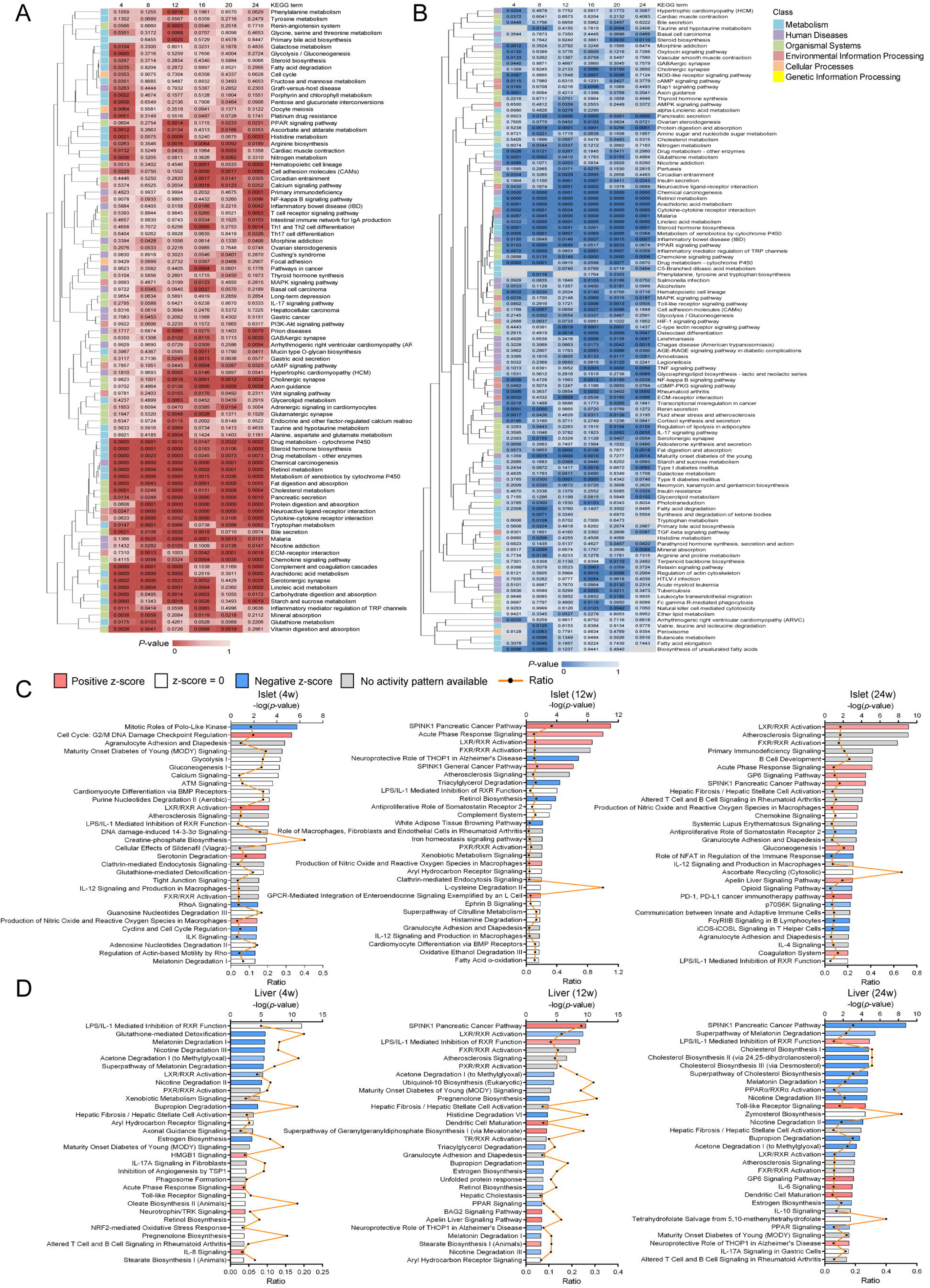
Temporal pathway enrichment analyses of DEGs. **(A, B)** Clustered heatmaps of islets (A) and liver (B) showing KEGG pathway enrichment analyses of DEGs across six time points. We displayed a pooled summary of the top 40 over-represented pathways at each time point in islets and liver. Clustering by columns was implemented using average linkage method. The intensity of color represents *P*-value. **(C, D)** IPA of islets (C) and liver (D) presenting the top 30 canonical pathways (y-axis) modulated by DEGs at week 4, 12 and 24 of diet. The significance of the association between dataset and canonical pathway was estimated by −log of *P*-value (Fisher’s exact test, top x-axis) and the ratio of genes that map to each canonical pathway (bottom x-axis).

### 3.6 Weight gene co-expression network analyses presenting trait-correlated modules

WGCNA algorithm was used in islets and liver to define trends in gene co-expression and look for a consensus network of genes that were correlated in all conditions. In total, we detected 42 co-expression modules in islets and 65 co-expression modules in liver (**Supplemental Table 3**). To identify gene clusters associated with traits of islet and liver function, together with other metabolic characteristics, we used linear regression models treating the module eigengenes (MEs) as dependent variables and the traits as independent variables. The individual metabolic profile was retrieved from the same mice subjected to RNA-sequencing. Those module-traits relationships were further used to select biologically meaningful modules for downstream analyses (**Supplemental Table 4**).

In total, eight islet modules were found to have a significant correlation with any one of the insulin secretion parameters calculated from IPGTT experiments. Genes from these modules were pooled and further analyzed by KEGG as shown in the top 30 annotated pathways. We observed enrichment of proliferative genes (*Cdc20*, *Bub1*, *Ccna2*, *Ccnb2* and *Cdk1*) in salmon module and energy metabolism related genes (*Atp5c1*, *Ndufs4*, *Cox5b*, *Cox7b* and *Ppa2*) in lightcyan1 and darkgrey modules (**Supplemental Figure 7A**). Notably, patterns of liver module expression across various traits exhibited strong and negative correlation with islet function, which might be attributed to liver as a crucial organ regulating systemic metabolism and glucose homeostasis. In liver, modules of interest were defined as those significantly correlated with more than 5 insulin secretion parameters. Among the 26 modules of interest, KEGG pathway enrichment suggested that brown and blue modules were evidently related to immune and inflammatory responses, including TNF signaling pathway (*Junb*, *Pik3cd* and *Ccdc88b*), toll-like receptor signaling pathway (*Tlr6*, *Ccl3* and *Ticam2*), T cell receptor signaling pathway and B cell receptor signaling pathway (*Jun*, *Fos*, *Cd22*, *Blnk* and *Vav1*) (**Supplemental Figure 7B**).

### 3.7 Proliferative and immune response signaling pathways underlying the crosstalk between islets and liver

Following the single-tissue, single-platform analyses of individual dataset, we considered the inter-tissue, multi-platform analyses using WGCNA and IPA by combining the islet and liver transcriptomic data together. To evaluate co-ordinated molecular interactions by identifying genes of islets and liver that were highly correlated, we generated a massive matrix of cross-tissue Pearson correlation coefficients for each gene of the islet modules to each gene of the liver modules in WGCNA. By taking the gene connectivity (membership) within its own module into consideration, combined correlation coefficients for each pair of islet and liver modules were calculated. In order to search for potential molecules underlying the inter-tissue communication contributing to islet dysfunction, we further looked into pairs of biologically meaningful modules, including 8 islets and 26 liver modules that were in association with insulin secretion traits. Among 208 (8×26) pairs, 8 pairs were identified with Pearson coefficient >0.4 and 17 pairs showed coefficient >0.35 and ≤0.4 (**Figure 6A**). We also detected “key genes” within a given module by calculating the gene’s correlation to a partner module. Within 8 pairs of islet and liver modules (Pearson coefficient >0.4), 34 islet and 25 liver “key genes” (**Supplemental Table 5**) in total that were highly representative of their own modules (membership >0.9) and highly correlated to a counterpart partner module (correlation>0.5) were found (examples of correlation between 4 pairs of modules as shown in **Figure 6B**). In islets, 23 “key genes” were differentially expressed between HFD and CD, which included genes encoding immune-related molecules (*Cxcr4* and *Fosb*) and transcription factors (*Egr1*). Early growth response-1 (EGR-1) was found to attenuate palmitic acid-induced endoplasmic reticulum stress and apoptosis in β-cells [32]. Among 17 differentially expressed “key genes” in liver, it is noticeable that C-X3-C motif chemokine receptor 1 (CX3CR1) can regulate β-cell function and impact insulin secretion through protection against the adverse effects of cytokines [33,34].

**Figure 6:**
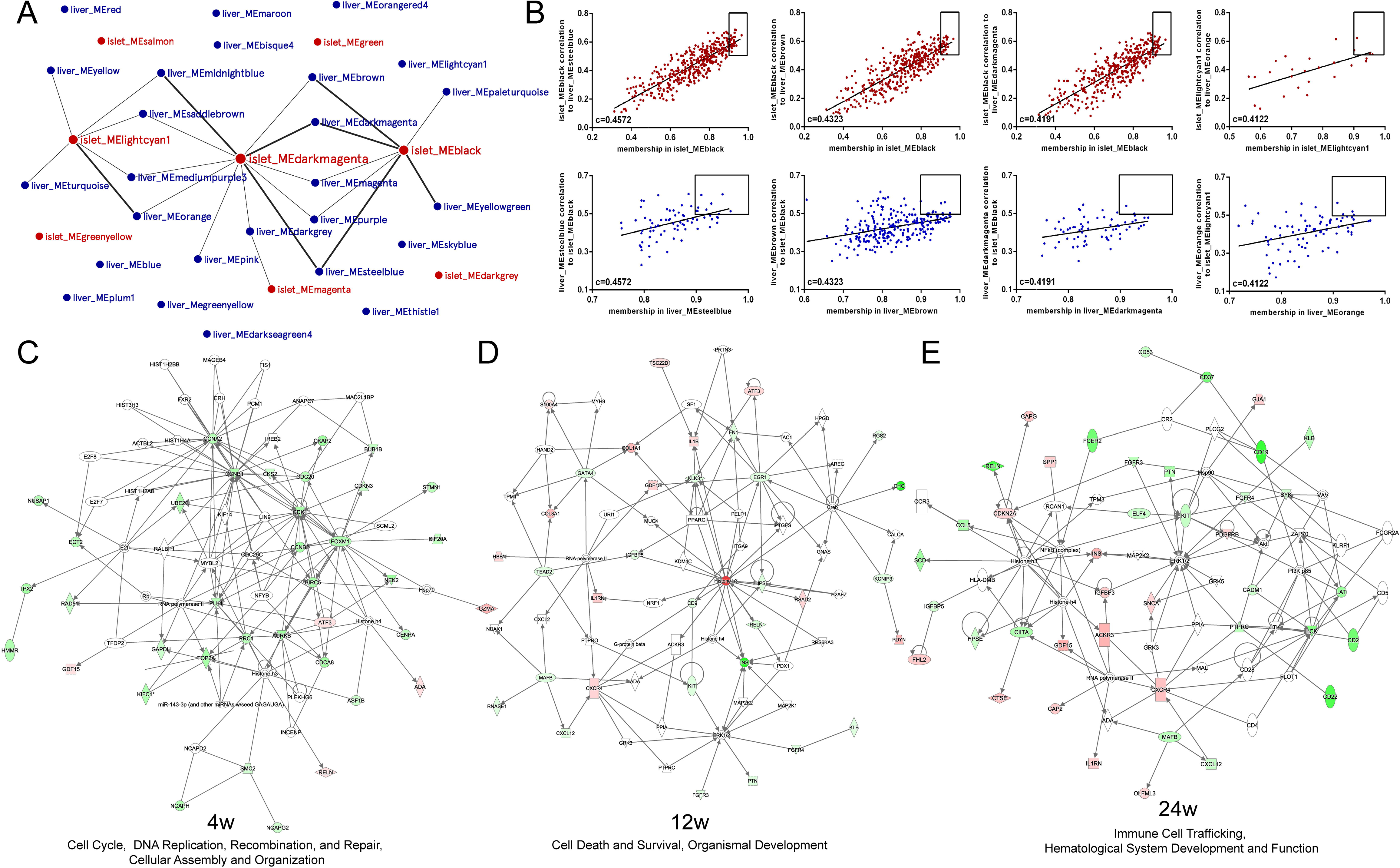
Key genes and signaling pathways involved in the crosstalk between islets and liver. **(A)** Network of trait-correlated islet and liver modules. In total, 8 islet (red circles) and 26 liver (blue circles) modules were defined as associated with insulin secretion traits. Among these 208 (8×26) pairs, 8 pairs of highly correlated modules were identified with Pearson coefficient >0.4 (linked with thick lines) and 17 pairs showed coefficient >0.35 and ≤0.4 (linked with thin lines). **(B)** Examples of key gene identification between two highly related trait-correlated islet and liver modules (Pearson coefficient >0.4). Each dot represents one gene and “key genes” (the top right of each graph) are those most correlated to a counterpart partner module (correlation >0.5) and also displayed a high module connectivity (membership >0.9). **(C, D, E)** Ingenuity network analyses using islet DEGs as the basis, and sequentially adding liver DEGs encoded for proteins that can be secreted at the same time point. The most dominant networks of week 4 (C), 12 (D) and 24 (E) were depicted. The up-regulated genes are in red and the down-regulated are in green.

To relate islet transcriptomic changes with liver perturbations, IPA was also applied for generating networks and identifying major integrative hubs. We postulated that one of the mechanisms underlying the islets-liver crosstalk might entail the direct effect of liver-derived secretome on islets. Consequently, we constructed networks for each time point by using the islet transcriptomic dataset as the basis, and sequentially adding the liver genes which were differentially expressed and encoded for proteins that can be secreted. Top scored networks showed potential interactions between several genes and secreted molecules implicated in cell cycle, DNA replication, recombination and repair during week 4 (**Figure 6C**), cell death and survival, organismal development around week 12 (**Figure 6D**), and immune cell trafficking, hematological system development and function at week 24 (**Figure 6E**) (**Supplemental Table 6**). Embedding the liver dataset allowed the identification of possible interconnected molecules, such as growth differentiation factor 15 (GDF15), activating transcription factor 3 (ATF3) and secreted protein acidic and rich in cysteine (SPARC).

### 3.8 The imbalance of T cell subpopulations in the development of diet-induced diabetes

Our pathway enrichment analyses of islets not only demonstrated a tight interplay between glucose metabolism and innate immune response, but also raised the intriguing possibility that adaptive immune response might be involved in the development of HFD-induced diabetes. In order to validate this postulation, we used flow cytometry to verify the temporal frequencies of T cell subpopulations both in spleens and draining lymph nodes during 24 weeks of feeding (**Figure 7A**). The proportion of splenic and lymphonodular T helper 1 (Th1) cells were markly increased in HFD mice compared with control mice since week 8 (**Figure 7B**). The disequilibrium of T helper 17 (Th17) cells also exhibited similar tendency as Th1 cells with elevated frequencies (**Figure 7D**), while the percentage of T helper 2 (Th2) cells were comparable between two groups (**Figure 7C**). Interestingly, a significant reduction of regulatory T cells (Tregs) during the early stage of obesity progression was also found (**Figure 7E**), whereas no difference was detected with regard to T follicular regulatory (Tfr) cells (**Figure 7F**). Alternations of T cell compartment exhibited a priming stage of decrement in anti-inflammatory Tregs at week 4−8, and an amplification stage of increase in proinflammatory Th1 and Th17 cells at week 12−24.

**Figure 7:**
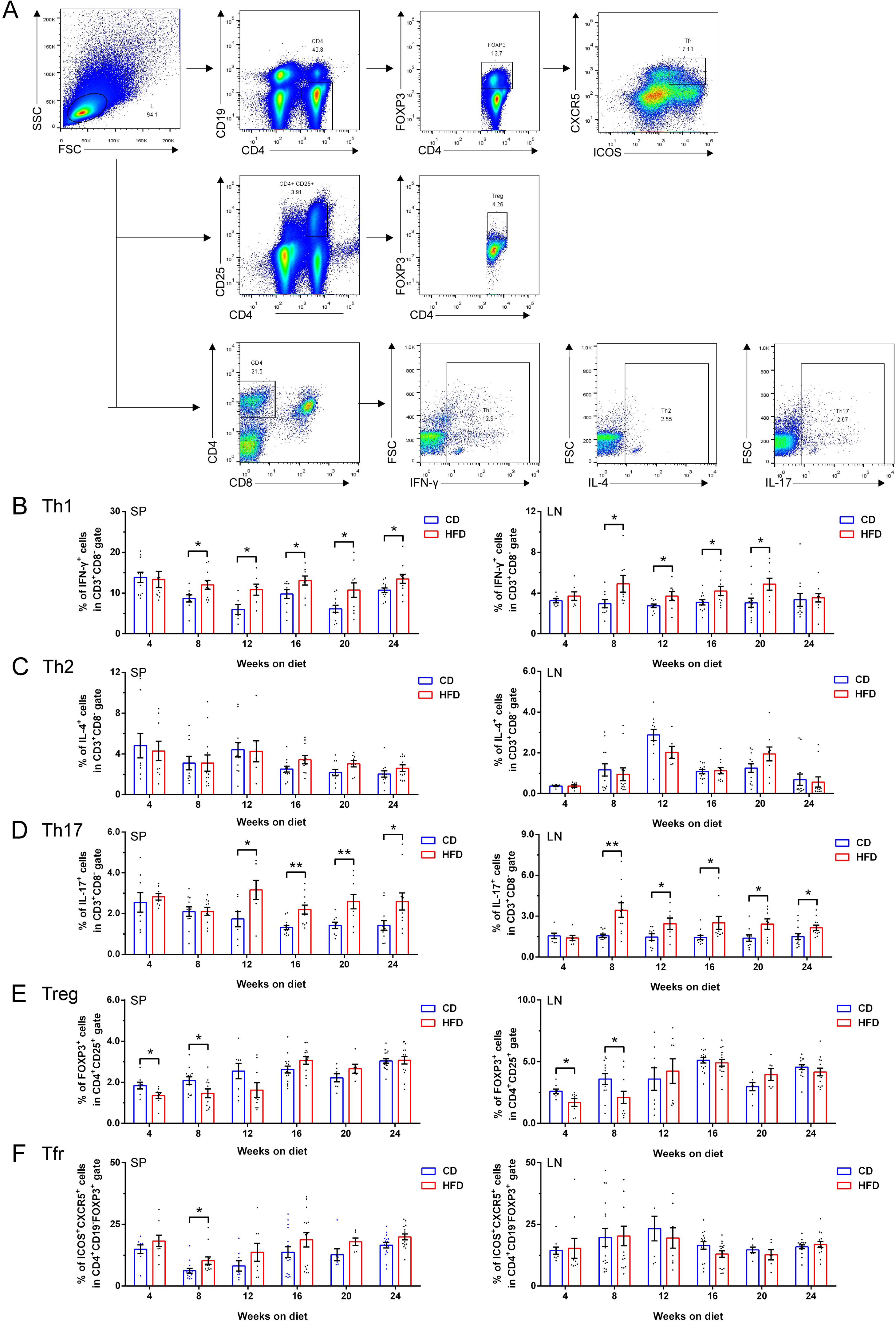
Charaterization of splenic and lymphonodular T cell subpopulations during 24 weeks of feeding. **(A)** Representative dot plots showing the gating strategies. **(B, C, D, E, F)** The frequencies of splenic (left column of scatter dot plots) and lymphonodular (right column of scatter dot plots) CD3^+^CD8^-^IFN-γ^+^ Th1 cells (B), CD3^+^CD8^-^IL-4^+^ Th2 cells (C), CD3^+^CD8^-^IL-17^+^ Th17 cells (D), CD4^+^CD25^+^FOXP3^+^ Tregs (E) and CD4^+^CD19^-^FOXP3^+^ICOS^+^CXCR5^+^ Tfr cells (F) (N≥7 mice/group). All data are expressed as mean±SEM and analyzed using unpaired two-tailed *t*-test. \**P*<0.05, \*\**P*<0.01.

## 4. DISCUSSION

Conflicting evidence in diabetic rodent models of whether hyperinsulinemia or insulin resistance is the primary driver indicates a lack of knowledge in the timeline and molecular basis after initiation of nutritional excess. The discrepancy might be attributed to confounding factors including genetic differences among strains, sex, age of diet onset and its duration, and diet composition. In the current study, we sought to avoid these variables by using the C57BL/6N mice with extended span. C57BL/6N mice do not harbor the mutation of nicotinamide nucleotide transhydrogenase (NNT) which has been associated with impaired β-cell function and glucose intolerance in some studies [35–37].

We identified that mild fed hyperglycemia was maintained in HFD mice throughout 24 weeks of diet and gradually alleviated by an ordered succession of adaptive β-cell response. Around week 4, fasting hyperinsulinemia first occurred in the absence of detectable fasting glucose elevation and impaired systemic insulin sensitivity. Such apparent “uncoupling” might indicate hyperinsulinemia is the primary effect caused by HFD feeding through direct stimulation of islet β-cells. This observation was also reported by several rodent studies focusing on very early time points of overfeeding [38–41]. Potential mediators of enhanced insulin secretion might be hyperlipidemia caused by nutrient overload as shown in elevated HDL-C and LDL-C at week 4. A plausible explanation for the glucose intolerance observed at week 4 despite of hyperinsulinemia and relatively intact insulin sensitivity is probably due to disturbed pulsatile pattern of insulin secretion in HFD mice. Changes in β-cell pulsatility could impact the efficacy of secreted insulin on its targets, particularly suppressing hepatic glucose production [42,43] and enhancing uptakes by peripheral tissues [44]. These impacts, however, could not be detected by HI/EG clamps with constant infusion of insulin and western blot of targeted tissues following an intraperitoneal insulin injection at week 4. From week 4−16, HFD mice exhibited progressively exaggerated insulin secretion in tandem with evident hyperglycemia. Meanwhile, defective tissue-specific insulin responsiveness showed variable time course with liver and skeletal muscle developing insulin resistance from week 16, while adipose tissue starting from week 8. After week 12, we observed robust immune and inflammatory response both in spleen and draining lymph nodes, which could further aggravate systemic insulin resistance. Interestingly, glucose intolerance was slightly improved as indicated by glycemic levels during IPGTT conducted around week 16−24. This phenomenon might be due to a compensatory increase in β-cell mass, and the subsequent hyperinsulinemia can fully compensate or even overtake the adverse effect of insulin resistance in maintaining glucose homeostasis. A major technical problem in assessing the roles of these two aspects in established obesity is that the measurement of each parameter may not be sufficiently precise and sensitive to dissect a causal sequence. Additionally, it is worthwhile noticing that although our data pointed towards the hypothesis that primary hyperinsulinemia initially causes insulin resistance, both mechanisms are not mutually exclusive and probably act in parallel at later stage.

Regardless of whether the trigger is over-nutrition, insulin resistance, or both, the transition from adaptive β-cell response to pathological β-cell response represents a crucial step in the development of obesity-induced diabetes. We concluded the greatest enhancement in β-cell proliferation was observed at week 4 on HFD, whereas a continuous increase in β-cell area was only apparent until week 8−12. This seemingly unexpected delay also takes place in maternal islets during pregnancy [45], when highly productive β-cells have to divide and accumulate before identifiable enlargement. We also noticed despite of the comparable proliferation rate between two diet regimens during week 12−24, a gradual growth in β-cell area was still detectable even at week 24, probably owing to the larger base of potentially proliferative β-cells in HFD mice.

Despite of compensation illustrated above, T2DM is distinctively characterized by a progressive reduction in β-cell function preceding the onset of overt disease. Our secretion data in vitro displayed a hypothesized temporal transition of HFD islets at three stages. Initially, enhanced β-cell function compensated insulin resistance by higher insulin release without significant β-cell mass increase. During the second stage, β-cell hyposensitivity to glucose and PA started to appear alongside with elevated basal secretion. Adaptive β-cell response, in combination with mass expansion, contributed to the rising plasma insulin concentration. Subsequently at the third stage, insufficient β-cell function was shown by decreased basal release and defective response to stimuli despite the ongoing β-cell mass increment. This supported the perspective which was also validated by alternations in β-cell calcium dynamics [46]:β-cell dysfunction, not islet mass decline, is the fundamental mechanism that fails at the initial stage of diet-induced diabetes.

In order to prevent β-cells from decline in function, it is important to investigate the underlying molecular mechanisms triggered by HFD involved in the progression toward T2DM. Identifying whether these pathways are protective/adaptive or deleterious/maladaptive would be helpful in developing strategies to reverse islet dysfunction [47]. Our transcriptomic data of islets and liver suggested several sequential events during shift from compensation to decompensation. Inter-tissue, multi-platform analyses showed that potential interactions of genes were implicated in cell cycle during week 4, organismal development around week 12, and immune cell trafficking at week 24. Beside HGF and SerpinB1 that have been already reported, we also found several genes that are noteworthy. GDF15, as an important signal in response to nutrition stress induced by long-term high-fat feeding, has been identified recently to improve insulin sensitivity [48,49] and protect against cytokine-induced β-cell apoptosis [50]. We observed an up-regulation of *Gdf15* in liver throughout 24 weeks of feeding, and identified it in the top scored IPA networks of all time points as an interconnected gene with islets (**Figure 6C, D and E**). Other potential molecules such as ATF3 and SPARC might be also involved in the inter-tissue crosstalk. Hypothalamus ATF3 could regulate systemic metabolism by altering insulin sensitivity and energy expenditure [51], while pancreatic ATF3 could modulate glucagon gene transcription under the circumstance of low glucose [52]. SPARC, primarily secreted by pancreatic stellate cells, could mediate the communication between stromal cells and endocrine cells by regulating β-cell survival [53]. However, the roles of ATF3 and SPARC in liver linking to islet function still remain elusive.

In conclusion, our study depicts a comprehensive landscape of temporal changes in islet and liver gene expression together with metabolic characteristic in HFD mice for 24 weeks. HFD mice exhibited progressively impaired glucose homeostasis with evident hyperinsulinemia and first-phase insulin secretion defect after week 4. Insulin, glucagon and somatostatin secretion in response to glucose or co-stimulated PA demonstrated a gradually deteriorated conversion from islet dysfunction to failure. HFD islet morphology showed increased abundance of β-cells whose proliferation peaked at week 4, with concomitant reduction in δ-cell and α-cell proportion. Ultrastructure of β-cell also presented decreased docked granules and deranged cristae of mitochondria. We identified impaired systemic insulin sensitivity from week 12 with variable time course in tissue-specific insulin action. Our islet and liver RNA-sequencing datasets outlined the impact of HFD on dynamics of molecular network at different stages. Correlation analyses of islet and liver modules with metabolic phenotypes illustrated that these two tissues jointly program β-cell adaption to irreversible impairment via cell cycle during week 4, organismal development around week 12, and immune cell trafficking at week 24. Alternations of T cell subpopulations validated the participation of adaptive immune response through a priming stage of decrement in anti-inflammatory Tregs, and an amplification stage of increase in proinflammatory Th1 and Th17 cells in diabetic progression. Future in-depth research of individual gene will help to discover potential diagnostic and therapeutic targets for human T2DM.

## AUTHOR CONTRIBUTIONS

Conceptualization, R.G., Q.F. and T.Y.; Methodology, R.G., Q.F., Q.Z. and T.Y.; Software, R.G. and Q.F.; Validation, R.G., Q.F., H.M.J. and Y.Q.; Formal analysis, R.G.; Investigation, R.G., Q.F., H.M.J., M.S., R.L.Z., Y.Q. and Y.Q.H.; Resources, K.F.X., X.Y.X. and H.C.; Data curation, R.G. and Q.F.; Writing (original draft preparation), R.G., Q.F., Q.Z. and T.Y.; Writing (review and editing), R.G., Q.F., H.M.J., M.S., R.L.Z., Y.Q., Y.Q.H., K.F.X., X.Y.X. and H.C; Visualization, R.G., Q.F., Q.Z. and T.Y.; Supervision, Q.Z. and T.Y.; Project Administration, T.Y. Funding Acquisition, Q.Z. and T.Y.

## ACKNOWLEDGEMENTS

This study was supported by National Natural Science Foundation of China (81830023), Postgraduate Research & Practice Innovation Program of Jiangsu Province (KYCX18_1474), scholarships from the China Scholarship Council (201908320471).

## Supplemental information inventory

**Supplemental Figure 1: Fat distribution, biochemical tests and glucagon levels in CD and HFD mice.**

****Supplemental Figure 2: Impaired liver structure, enlarged islet mass and altered ultrastructure in**α**-cells and**δ**-cells of HFD mice.****

**Supplemental Figure 3: Longitudinal assessment of tissue-specific insulin sensitivity in quadriceps femoris, gastrocnemius and white adipose tissue.**

****Supplemental Figure 4: PCR validation of transcriptomic data and schematic of time course in HFD mice.****

**Supplemental Figure 5: Time course expression of factors regulating endocrine pancreas development and **β**-cell mass.**

****Supplemental Figure 6: Time course expression of pathways involved in hepatic insulin resistance.****

**Supplemental Figure 7: Pathway enrichment analyses of trait-correlated modules.**

**Supplemental Table 1: Summary results of sequencing reads per sample.**

**Supplemental Table 2: Primer sequences of quantitative RT-PCR for validation.**

**Supplemental Table 3: List of islet and liver modules.**

**Supplemental Table 4: (A) Heatmap showing the correlation matrix between islet modules and phenotypic traits. (B) Heatmap showing the correlation matrix between liver modules and phenotypic traits.**

****Supplemental Table 5: (A) List of islet key genes found highly related to trait-correlated liver modules. (B) List of liver key genes found highly related to trait-correlated islet modules.****

**Supplemental Table 6: (A) Ingenuity network analysis of crosstalk between islets and liver at 4 weeks of diet. (B) Ingenuity network analysis of crosstalk between islets and liver at 12 weeks of diet. (C) Ingenuity network analysis of crosstalk between islets and liver at 24 weeks of diet.**

