## Supplemental Figures for "Temporal Metabolic Characteristics and Transcriptomic Landscape of Islets and Liver Reveal Dynamic Pathophysiology and Interorgan Crosstalk in High-fat Diet-induced Diabetes"

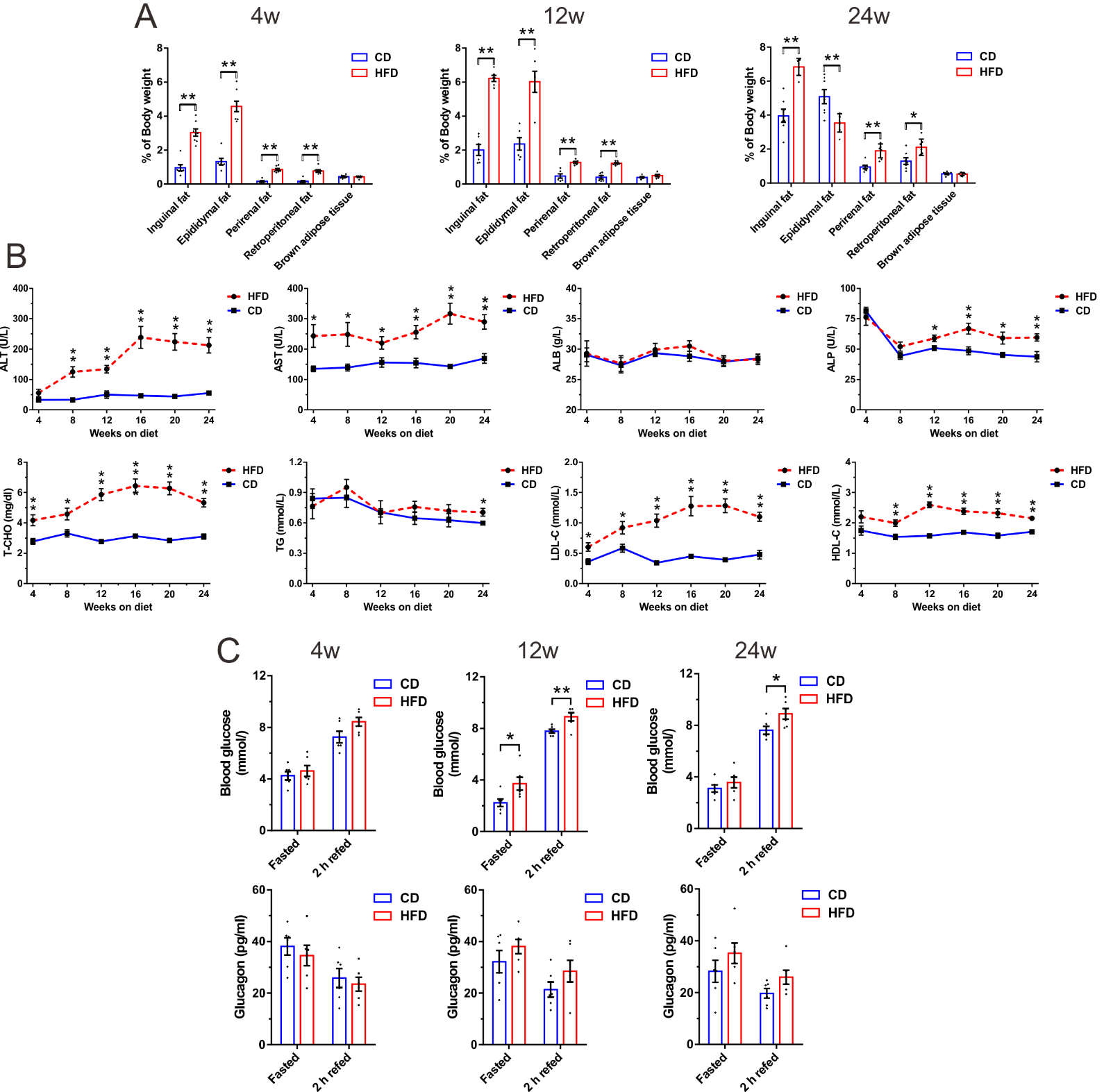

**Supplemental Figure 1: Fat distribution, biochemical tests and glucagon levels in CD and HFD mice. (A)** Analyses of inguinal, epididymal, perirenal, retroperitoneal fat depots and brown adipose tissue normalized to body weights (n=6−8 mice/group). **(B)** Biochemical tests of serum alanine aminotransferase (ALT), aspartate aminotransferase (AST), alkaline phosphatase (ALP), albumin (ALB), total cholesterol (T-CHO), triglyceride (TG), high-density lipoprotein cholesterol (HDL-C), and low-density lipoprotein cholesterol (LDL-C) after overnight fast (N=8 mice/group). **(C)** Blood glucose and plasma glucagon levels in fasted and 2 h refed HFD and CD mice after 4, 12 and 24 weeks of diet (N=6 mice/group). All data are expressed as mean±SEM and analyzed using unpaired two-tailed *t*-test. **P*<0.05, ***P*<0.01.

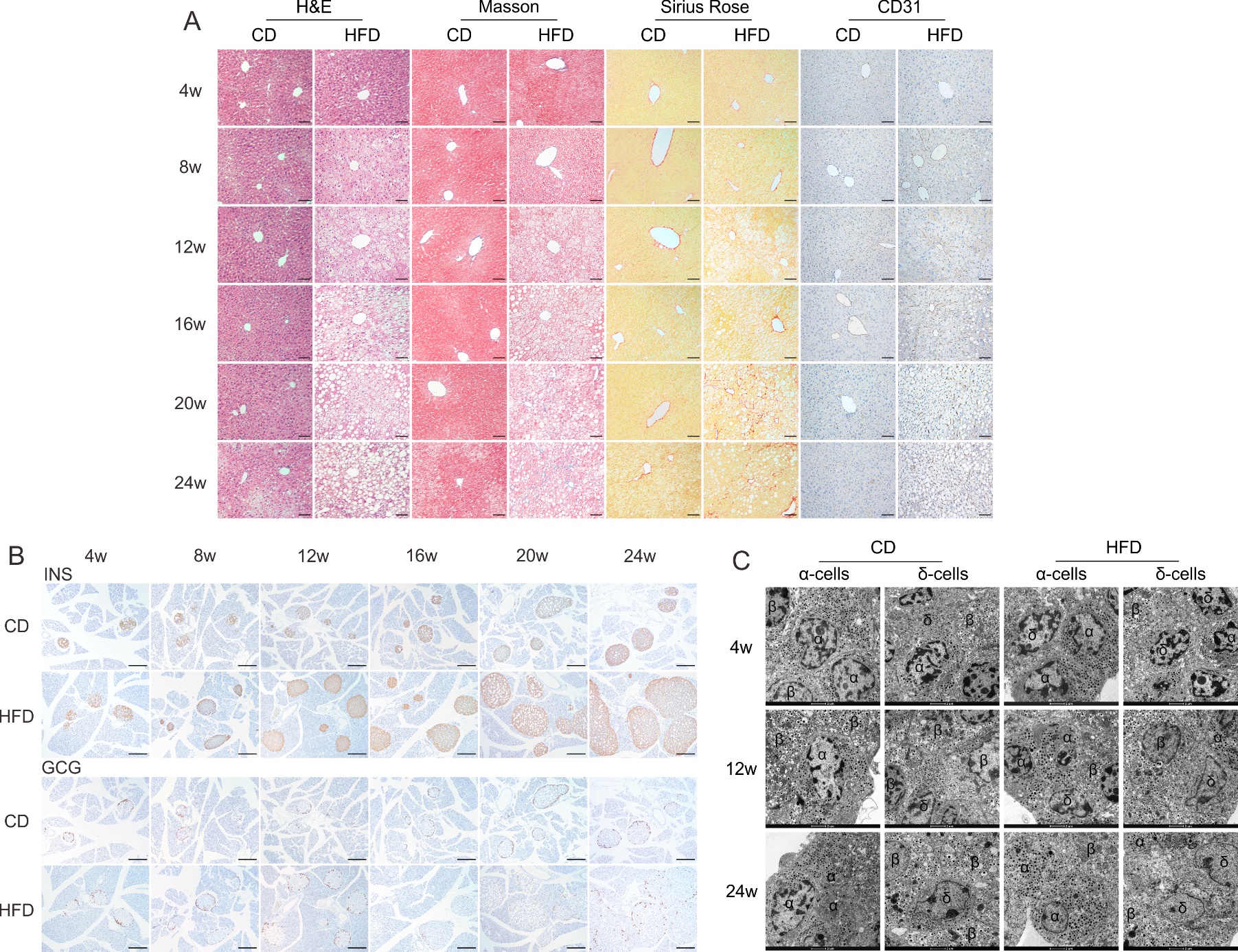

**Supplemental Figure 2: Impaired liver structure, enlarged islet mass and altered ultrastructure in α-cells and δ-cells of HFD mice.** **(A)** Representative photomicrographs of liver sections stained with H&E for liver structure, Masson’s Trichrome and Sirius Rose staining for collagen fiber deposition, and anti-CD31 for microvessel density. Images are representative of 4 mice per group (scale bar: 20 μm). **(B)** Representitive immunohistochemical images showing consecutive pancreatic sections labeled either for insulin or glucagon. Images are representative of 4 mice per group (scale bar: 50 μm). **(C)** Representitive electron micrographs showing ultrastructural changes in α-cells and δ-cells. Images are representative of 8−12 α-cells or δ-cells/6−8 islets/3 mice per group (scale bar: 2 μm). All data are expressed as mean±SEM and analyzed using unpaired two-tailed *t*-test. **P*<0.05, ***P*<0.01.

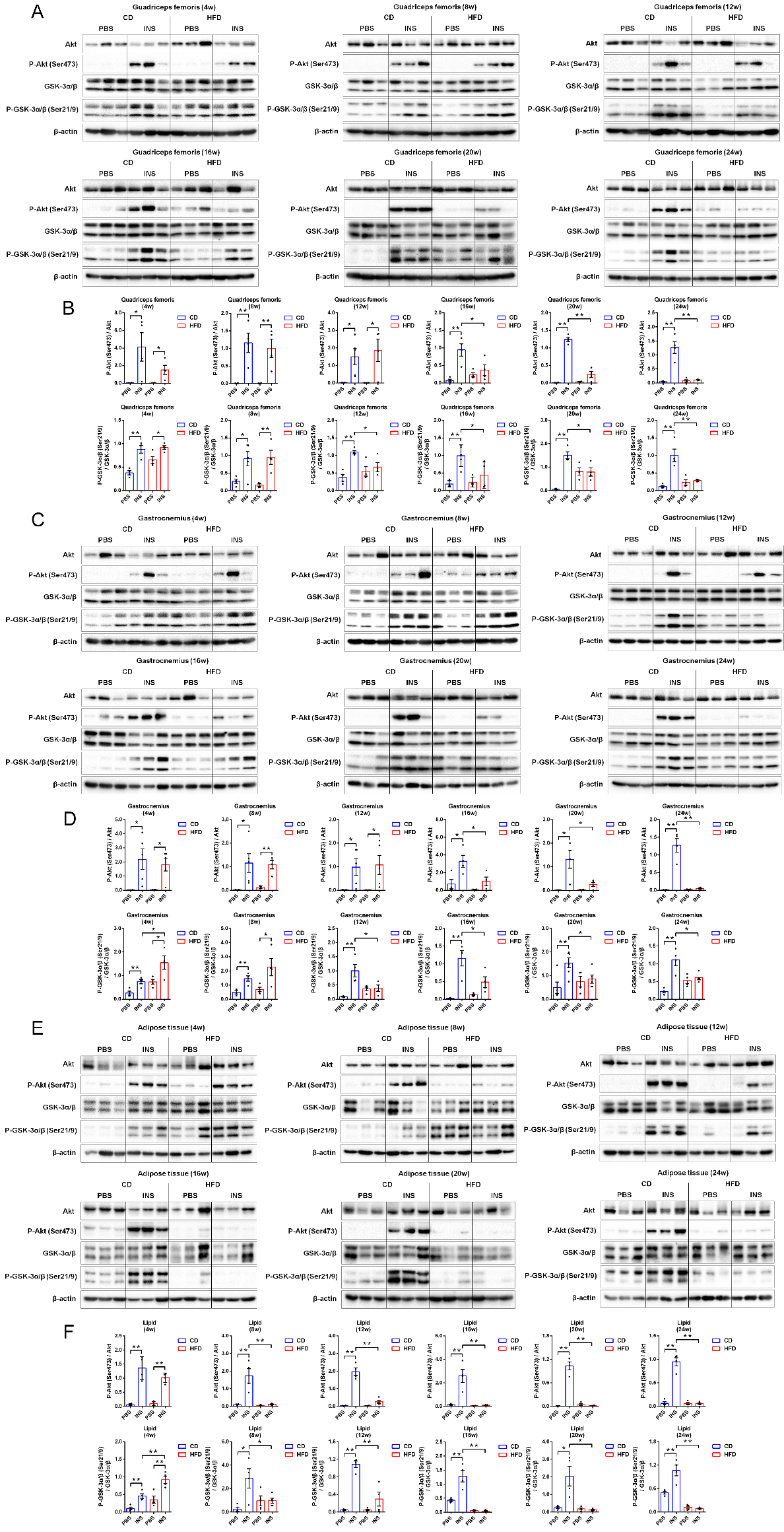
**Supplemental Figure 3: Longitudinal assessment of tissue-specific insulin sensitivity in quadriceps femoris, gastrocnemius and white adipose tissue.** **(A, B)** Immunoblot (A) and quantification (B) of Akt (Ser473) phosphorylation status relative to total Akt and GSK-3α/β (Ser21/9) phosphorylation status relative to total GSK-3α/β in quadriceps femoris (N=4−6 mice/group). **(C, D)** Immunoblot (C) and quantification (D) of these parameters in gastrocnemius (N=4−6 mice/group). **(E, F)** Immunoblot (E) and quantification (F) of these parameters in white adipose tissue (N=4−6 mice/group). All data are expressed as mean±SEM and analyzed using unpaired two-tailed *t*-test. **P*<0.05, ***P*<0.01.

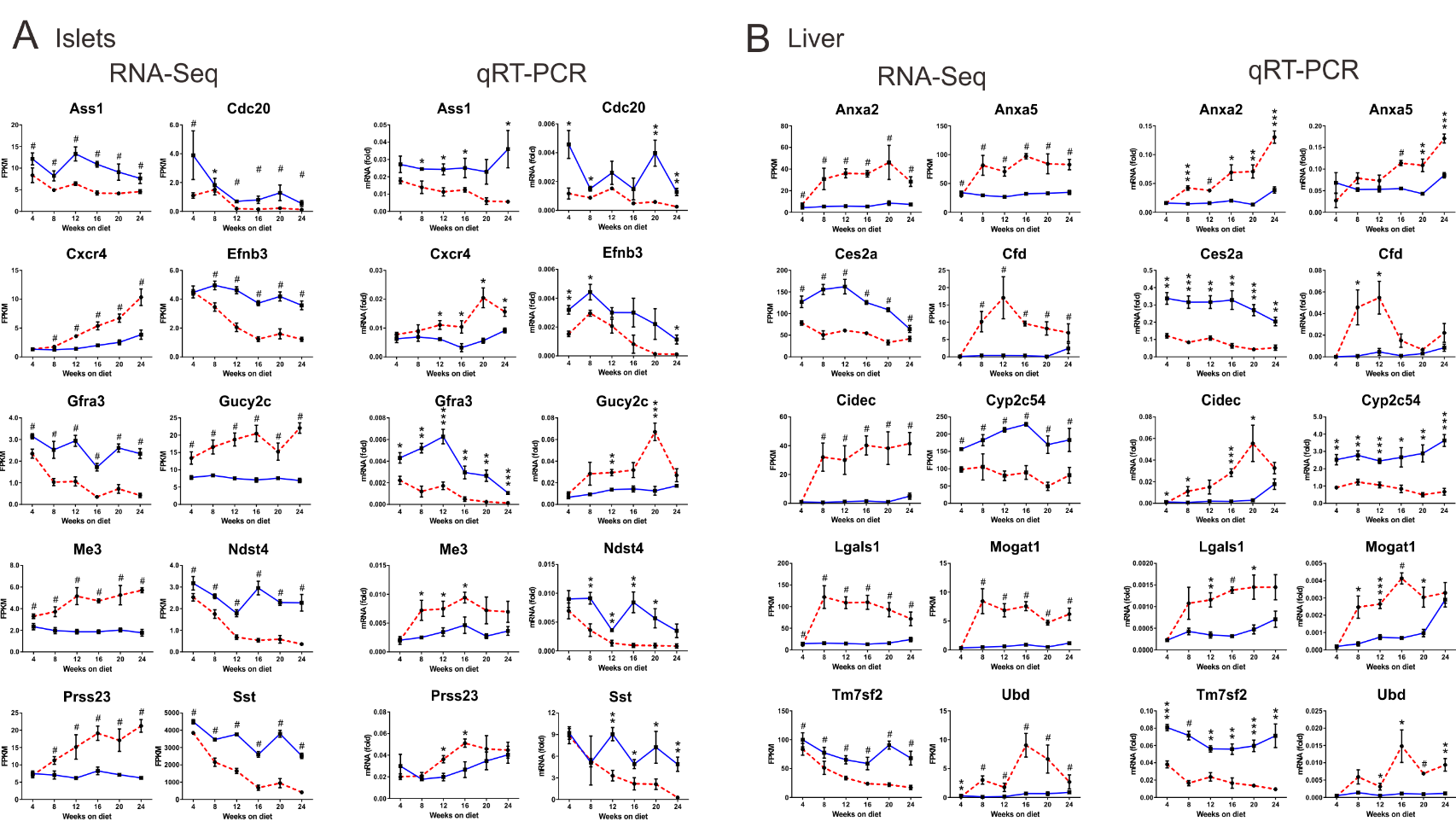

**Supplemental Figure 4: PCR validation of transcriptomic data and schematic of time course in HFD mice. (A, B)** Random selection of 10 differentially expressed genes for the validation of islet (A) and liver (B) transcriptomic data. The expression levels of these genes were measured in independent islet and liver samples (N=4 mice/group) (right column). The corresponding RNA-sequencing results of each gene were listed side by side for comparison (left column). All data are expressed as mean±SEM. Left column: *adjusted *P*<0.05, **adjusted *P*<0.01, ***adjusted *P*<0.001, ^#^adjusted *P*<0.0001. Right column: **P*<0.05, ***P*<0.01, ****P*<0.001, ^#^*P*<0.0001.

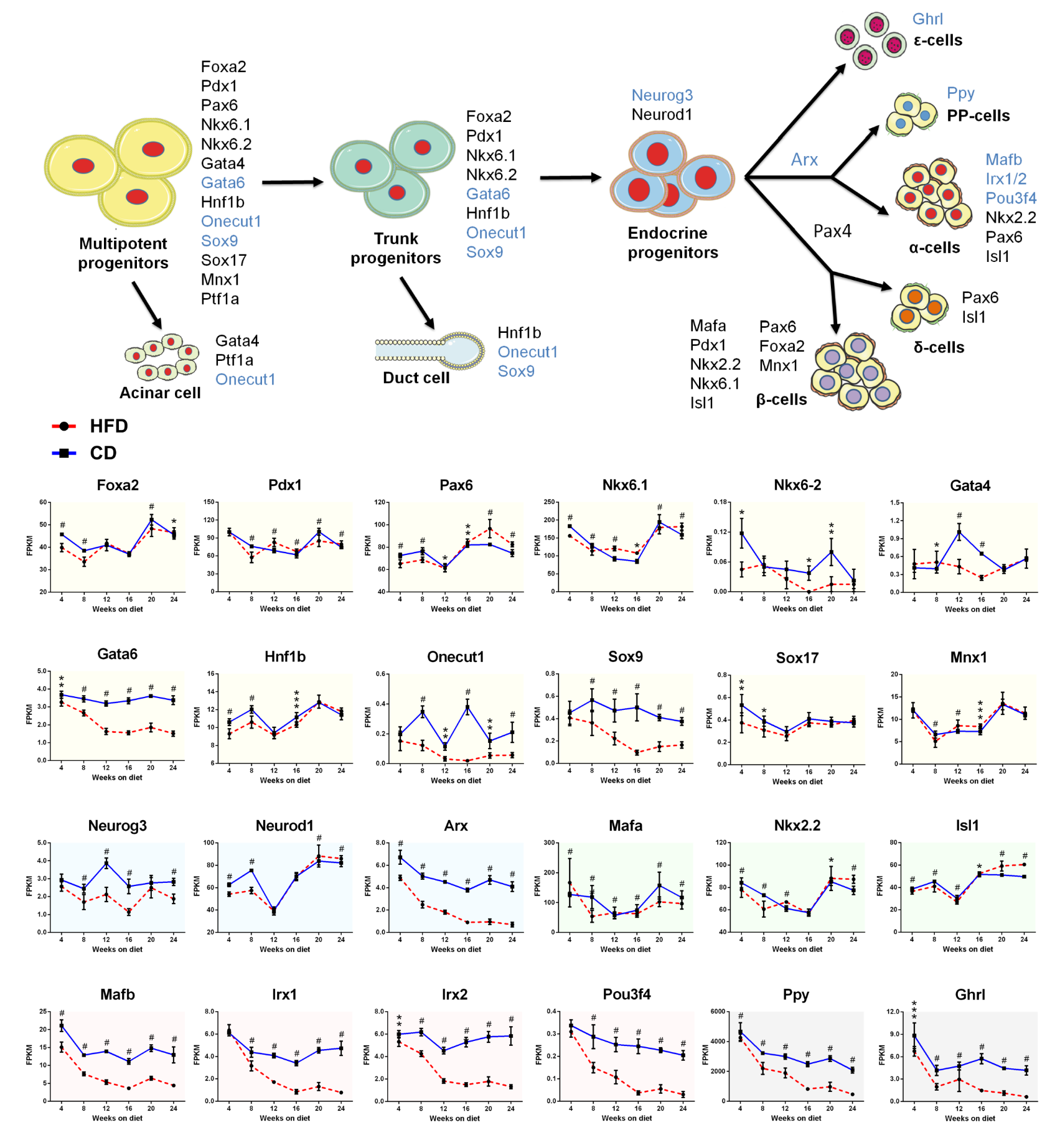
**Supplemental Figure 5:** **Time course expression of factors regulating endocrine pancreas development and β-cell mass.** The schematic representation depicted an overview of pancreatic progenitors toward differentiated lineages. Key transcription factors involved in each differentiation step are indicated and those significantly down-regulated are colored in blue. The time course expression data for these key genes are displayed in the line charts. Yellow, blue, green, red and grey shaded areas respectively refer to genes participating in the differentiation of multipotent progenitors, trunk progenitors, endocrine progenitors, hormone secreting α-, β-, δ-cells and PP-, ε-cells. All data are expressed as mean±SEM. *adjusted *P*<0.05, **adjusted *P*<0.01, ***adjusted *P*<0.001, ^#^adjusted *P*<0.0001.

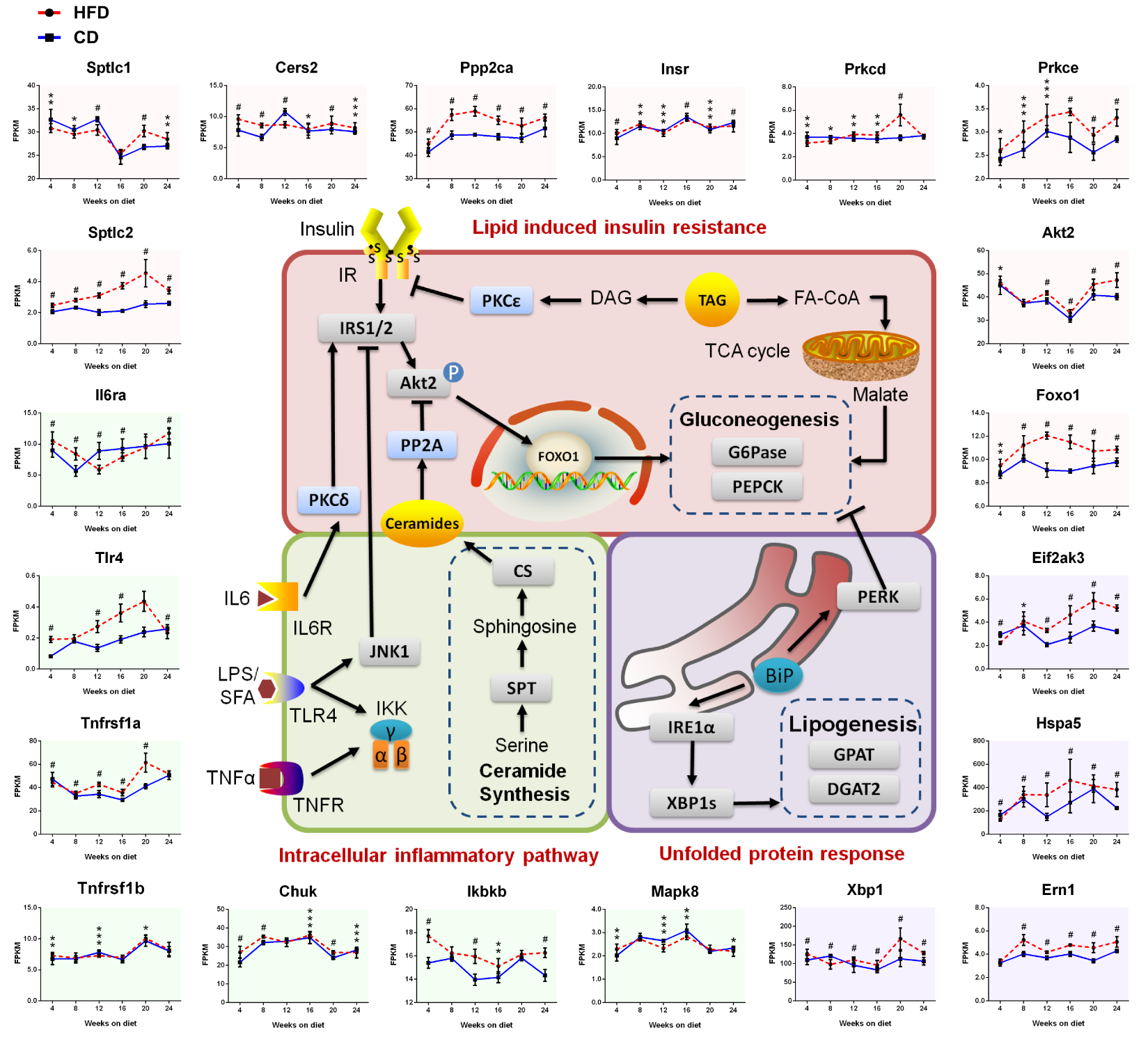
**Supplemental Figure 6: Time course expression of pathways involved in hepatic insulin resistance.** The schematic representation depicted three major mechanisms for hepatic insulin resistance. The time course expression data for these key genes are displayed in the line charts. Red, green and purple shaded areas respectively refer to genes participating in lipid-induced insulin resistance, intracellular inflammatory pathway and unfolded protein response. All data are expressed as mean±SEM. *adjusted *P*<0.05, **adjusted *P*<0.01, ***adjusted *P*<0.001, ^#^adjusted *P*<0.0001.

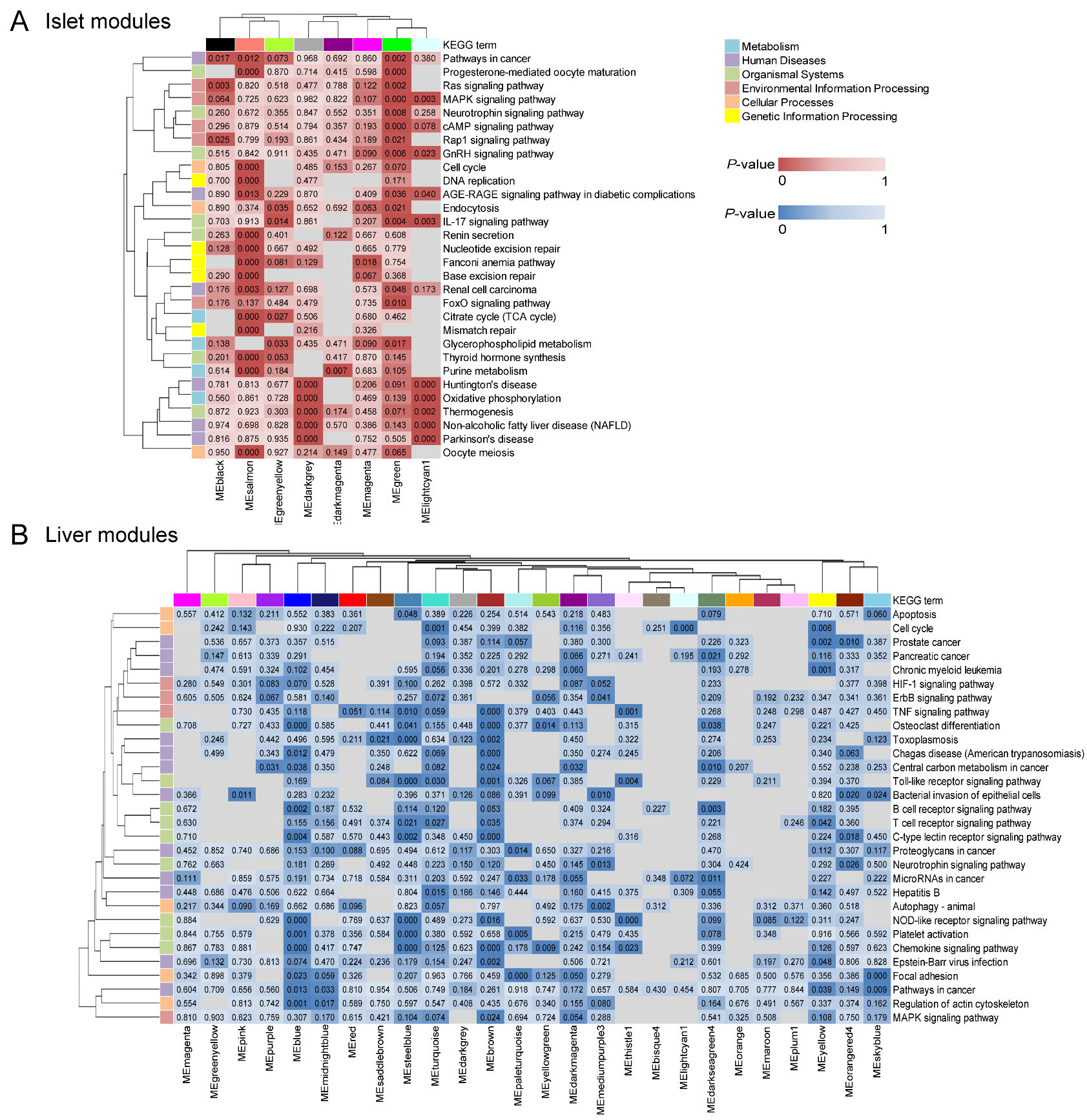

**Supplemental Figure 7:** **Pathway enrichment analyses of trait-correlated modules.** **(A, B)** Clustered heatmaps of islets (A) and liver (B) showing KEGG pathway enrichment analyses of trait-correlated modules. We displayed the overall top 30 over-represented pathways among these trait-correlated modules in islets and liver. Clustering by rows and columns was implemented using average linkage method. The intensity of color represents *P*-value.
